# Formation of the active Hermes transpososome is driven by asymmetric DNA binding of BED domains

**DOI:** 10.1101/2022.09.16.508221

**Authors:** Laurie Lannes, Christopher M. Furman, Alison B. Hickman, Fred Dyda

## Abstract

The cut-and-paste *Hermes* DNA transposase stands out among the transposases that have been biochemically or structurally characterized so far. Many transposases function as dimers, but the Hermes transposase forms a tetramer of dimers to achieve its active form *in vivo*. Intriguingly, the transposition complex, or transpososome, relies on only one dimer to perform the enzymatic reactions necessary to the mobilization of its transposon. Our investigation combining biochemical and structural approaches shows that the Hermes octamer extensively interacts with its transposon left-end (LE) engaging the BED domains of three Hermes protomers belonging to three dimers. By contrast, the right-end (RE) is entirely deprived of such interaction inside the transpososome. Our work suggests that formation of the *Hermes* synaptic complex is sequential and relies on the considerable difference of affinity of the transposase towards its transposon ends. Thus, we propose that Hermes dimers multimerize to gather enough BED domains to find the LE among the abundant genomic DNA, facilitating the subsequent interaction with the RE, most likely solely based on recognition of its terminal inverted repeat.

## Introduction

Transposable elements (TE) are discrete genomic regions that can move from one position to another within genomes. TEs have been found in all eukaryotic kingdoms as both active and inactive forms and can make up a large portion of eukaryotic genomes (Biémont and Vieira 2006). TEs have been a major force in the shaping of genomes and the evolution of species (Wells and Feschotte 2020).

Most of the eukaryotic class II or DNA transposons move by cut-and-paste transposition that consists in the excision of the transposon by generating double-stranded breaks (DSB) at the ends of the element and its reintegration elsewhere typically without specificity (Fig. 1A insert). An autonomous transposon is delimited by two ends typically arranged as terminal inverted repeats (TIRs) with one or more genes between them. One of these genes encodes the transposase, an enzyme that mobilizes the transposon by carrying out the necessary nuclease and trans-esterification activities. Cut-and-paste transposases contain a ribonuclease H-like (RNase H-like) catalytic domain in which a triad of carboxylate residues, arranged as DDE/D, coordinates two Mg^2+^ metal ions forming the active site (Yuan and Wessler 2011). The transposase typically assembles into a multimer that recognizes and brings together the TIRs by relying on site-specific DNA binding domain(s) such as helix-turn-helix or zinc-finger domains which can be either upstream or downstream of the RNase H-like catalytic domain (Feschotte and Pritham 2007). After synapse formation, DSBs are created at the ends of the TIRs, liberating the transposon, and the assembly finds its target DNA to integrate the TE.

**Figure 1:**
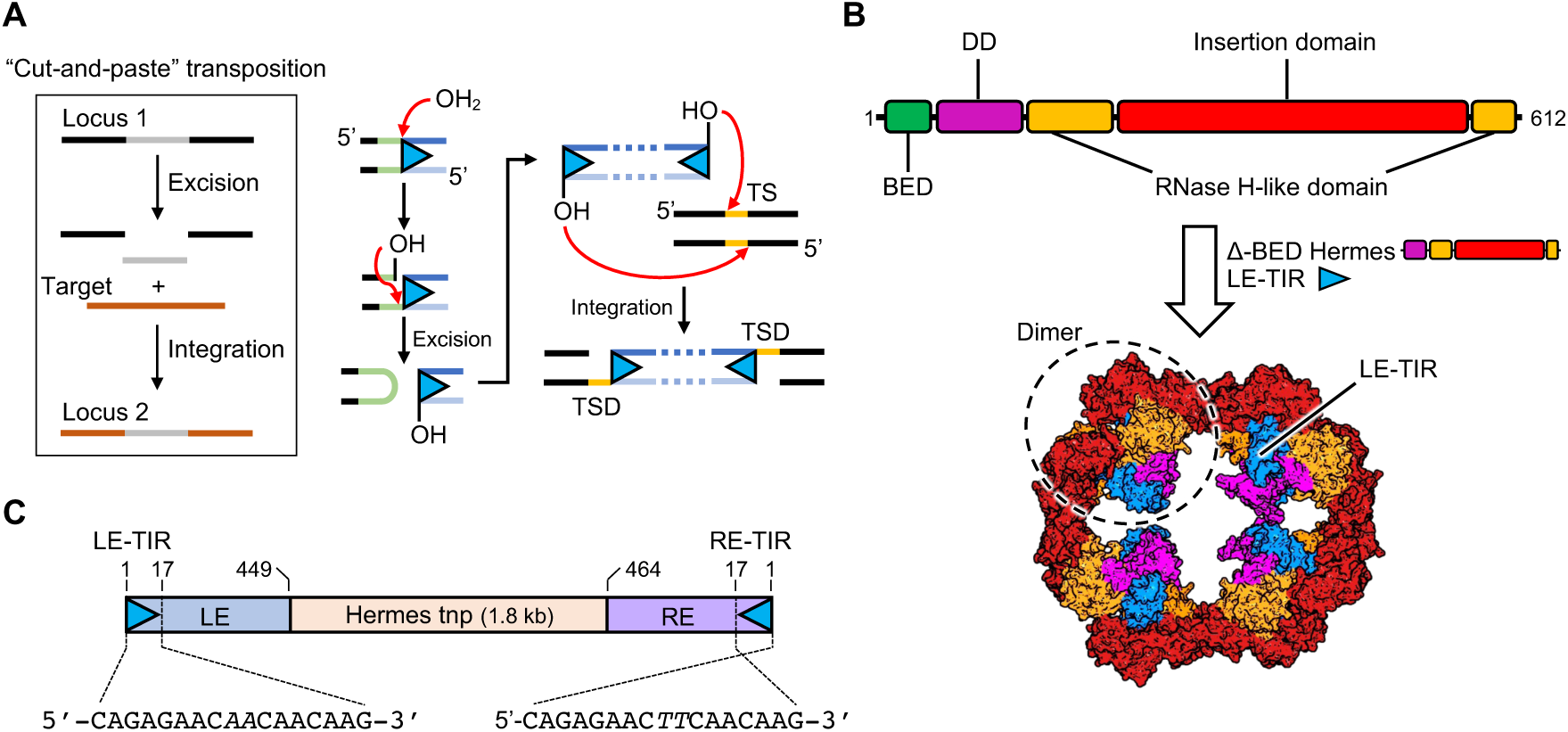
The eukaryotic *Hermes* transposon system. A) The “cut-and-paste” (excision/integration) transposition mechanism of the *Hermes* transposon and its chemistry. The red arrows describe nucleophilic attacks on phosphate groups. TS: target site. TSD: target site duplication. B) Top: schematic of the domain organization of the Hermes transposase. DD, dimerization domain. Bottom: octameric quaternary structure (tetramer of dimers) of BED-truncated (Δ-BED) Hermes in complex with the Left End Terminal Inverted Repeat (LE-TIR) DNA (PDB: 4D1Q, (Hickman et al. 2014)). C) Schematic of the *Hermes* DNA transposon. The transposon has a Left and Right End (LE and RE, respectively) capped by two imperfect Terminal Inverted Repeats (TIRs) that differ from one another by two swapped base pairs marked in italics. The ends flank the gene that encodes for the Hermes transposase (tnp).

The *hAT* superfamily named after its first members *hobo*, *Activator* (*Ac*) and *Tam3* is one of the largest DNA transposon superfamilies. *Hermes,* a *hAT* transposon isolated from the genome of the housefly *Musca domestica* has been characterized biochemically (Hickman et al. 2018; Zhou et al. 2004), and the three dimensional structure of its N-truncated form with its catalytic core (hereafter referred to as “Δ-BED”) has been solved (Fig. 1B) (Hickman et al. 2014). The *Hermes* transposon is ~3 kb and is delimited by two dissimilar ends of ~450 bp referred to as the left-end (LE) and the right-end (RE). The ends are capped by two imperfect 17 bp TIRs that differ from each other by two swapped base pairs (Fig. 1C). The Hermes transposase is composed of 612 amino acids and is organized in four domains (Fig. 1B).

The chemistry of *hAT* transposition is well known and differs from other cut-and-paste mechanisms by the formation of a hairpin on the strands flanking the transposon at the excision step (Fig. 1A) (Hickman and Dyda 2016; Zhou et al. 2004). The *hAT* transposases show insertion preference in 8 bp target sites with little specificity with the *Ac* family favoring 5’-nTnnnnAn sites (Evertts et al. 2007; Kim et al. 2011; Kondrychyn et al. 2009), and the *Buster* family preferring 5’-nnnTAnnn sites (Arensburger et al. 2011). Staggered integration across the target site generates 8 bp target site duplications (TSD) (Fig. 1A).

In principle, two transposase protomers, each carrying one catalytic domain with its active site, could assemble into a dimer to liberate two transposon ends and integrate them into a target site. In fact a number of DNA transposases from both the prokaryotic and eukaryotic domains appear to work as dimers (for example, Tn5, ISCth4, Mos1, the P element, and Transib; (Davies et al. 2000; Ghanim et al. 2019; Kosek et al. 2021; Liu, Yang, and Schatz 2019; Richardson et al. 2009) suggesting that a dimer is a minimal oligomerization unit (Davies et al. 2000). However, there are also examples of higher order oligomers. The bacteriophage MuA transposase forms a tetramer in its active form (Lavoie et al. 1991; Montaño, Pigli, and Rice 2012), whereas the related retroviral integrases can form even higher order oligomers, at least *in vitro* (Li et al. 2006; Passos et al. 2017).

Architecturally, the Hermes transposase is unusual as it spontaneously forms a ring-shaped tetramer of dimers (Fig. 1B). The dimers tightly associate using an “intertwined” dimerization domain (DD; Fig. 1B). The quaternary structure is held together by the swapping over of an α-helix from the insertion domain between adjacent dimers (Hickman et al. 2005, 2014). Deletion of the swapped helix prevents octamer formation, converting Hermes into dimers. Remarkably, these dimers are fully competent for all the enzymatic activities of Hermes *in vitro*; in fact, they are more active than wild-type Hermes. Yet they are inactive for transposition in cell-based assays (Hickman et al. 2014). The available biochemical and structural data provide no explanation for this substantial contradiction between *in vitro* and in cell experiments.

All *hAT* transposases are predicted to contain an N-terminal BED zinc-finger domain organized around a conserved CCHH or CCHC motif that coordinates Zn^2+^ (Aravind 2000), but no structural information is available for Hermes as the constructs used in previous structural work lacked the first 78 amino acids. Nevertheless, this domain is indispensable for the excision step *in vitro* (Hickman et al. 2014). We have previously suggested that the BED domain of the Hermes transposase recognizes short subterminal repeats (STRs) interspersed in its transposon ends (Hickman et al. 2014). Even though the transposition chemistry occurs at the very tips of TEs, several *hAT* transposons have DNA end requirements that span far beyond the TIRs. For instance, *Tol2* needs the terminal 200 and 150 bp of its LE and RE, respectively, for excision and transposition (Urasaki, Morvan, and Kawakami 2006); *Tag1* about 100 bp from both ends to generate comparable transposition rates as the full ends (Liu et al. 2001); and *Ac* at least 200 bp from both ends (Coupland et al. 1989). In the case of *Tol2*, it has been shown that mutation of STRs reduced the excision activity (Urasaki et al. 2006). Given the number of STRs in the *Hermes* ends, a dimer with only two BED domains does not appear sufficient if the binding of more than two is needed. Therefore, it is plausible to assume that interaction with multiple STRs is needed for in cell activity, and that a higher order assembly such as the octamer with its eight BED domains would be one way to supply a sufficient number of BED domains. However, the architectural features of such an assembly are not known.

As we suspected that deciphering the role of the BED domain in transposon end binding and understanding of how the transposase interacts with its ends could explain the striking discrepancy between *in vitro* and in cell results, we undertook a mechanistic study using biochemistry, X-ray crystallography, and cryo-electron microscopy (cryo-EM).

## Results

### The N-terminal domain of the Hermes transposase is a DNA binding domain

Although the first 78 amino acids of Hermes that include the BED domain are indispensable for transposition *in vivo* and cleavage *in vitro*, they do not directly participate in the recognition of the TIR or in the catalytic activity of the transposase (Hickman et al. 2014, 2018). Despite the critical role of the BED domain, its DNA binding site has not yet been established. We therefore first sought to confirm the DNA binding activity of Hermes’ BED domain and determine if such activity was specific. We expressed the region 1-78 of the Hermes transposase (hereafter “BED”) in *E. coli* and purified it (Fig. S1A) to perform interaction assays. While the LE and RE of the *Hermes* transposon are 449 bp and 464 bp, respectively (Warren, Atkinson, and O’brochta 1994), the proximity in sequence space of the BED domain to the part of Hermes with known three-dimensional structure suggests that DNA binding would likely occur towards the interior portion of the TIR. This notion is also supported by previous activity data using mutated Hermes LE (Hickman et al. 2014).

We used a double stranded DNA (dsDNA) oligonucleotide that spanned bp 11-27 of the LE (LE11-27, Fig. 2A) to probe the DNA binding of the BED domain. As a control, a randomized 17-mer (ran17) was used that had no common features with LE11-27. The DNAs were titrated with increasing amount of protein from one to three equivalents and analyzed by size exclusion chromatography (SEC). The resulting chromatograms are presented in Fig. 2A. Binding was assessed by the shift of the DNA peak, the shape of any resulting complex peak, and whether we observed free DNA or protein as a function of titration. Although ran17 showed evidence for only weak binding by BED, LE11-27 formed a tight and stable complex that saturated at a 1:2 ratio of DNA to protein.

**Figure 2:**
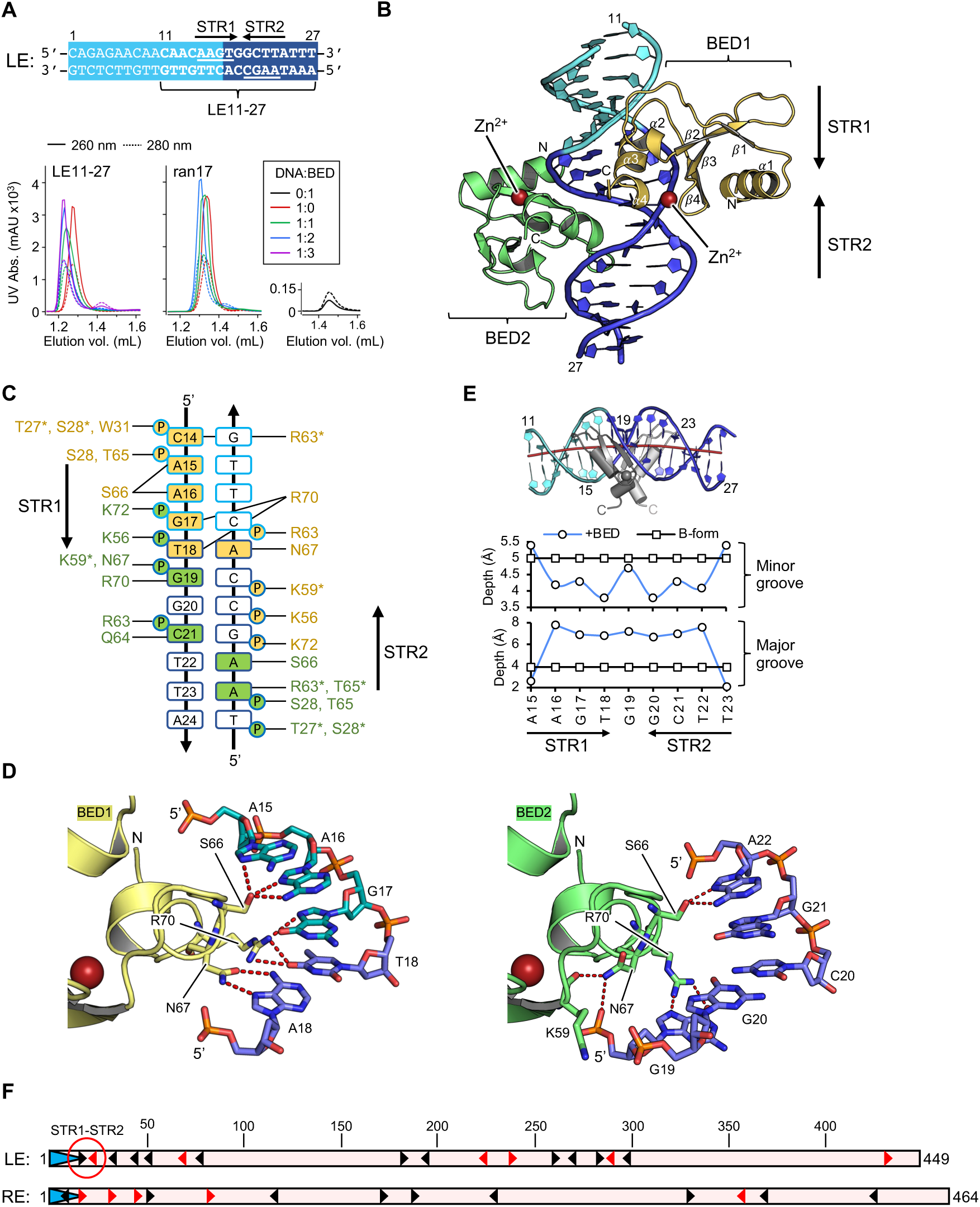
The N-terminal BED domain of the Hermes transposase. A) The sequence of the first 27 base pairs of the *Hermes* transposon Left End (LE). The Terminal Inverted Repeat (TIR) is in light blue while the subterminal region is in dark blue. The sequence in bold (LE11-27) was used as binding substrate for the BED domain. It presents a quasi-palindromic motif composed of two subterminal repeats (STRs), STR1 and STR2. The interaction was monitored by analytical size exclusion chromatography. The sequence of the control oligonucleotide (ran17) is 5’-CGCGATGAGTTCTCGAC-3’. The DNA-to-BED ratios of the samples are summarized in the insert. B) Co-crystal structure of two zinc-bound BED domains in complex with LE11-27 DNA. The DNA color code is the same as in A) and the top-strand is numbered. The N- and C-terminal ends of the proteins are indicated; and the secondary structure numbering is indicated on BED1 (in yellow). The Zn^2+^ ions are represented as red spheres. C) Summary of the protein-DNA hydrogen bond network in the crystal structure. The BED binding sites overlap with the STR1 and STR2 motifs. The amino acids in yellow and green belong to BED1 and BED2, respectively. The residues marked with a star interact via their main chain, while all the other engage their side chain. The top-strand of LE11-27 is numbered. The outline color of the bases follows the same code as in A), while the filling color identifies the BED domain interacting with them. Only the phosphate groups (P) involved in hydrogen bonds with the proteins are pictured. The sugar puckers are not represented. D) The hydrogen-bonding network between S66, N67 and R70 of the Hermes BED domain with the LE-STR1 (left) and STR2 (right) of the LE11-27 DNA. The H-bond network between the DNA and the protein is pictured by red dashed lines. The alternative conformations of N67 and R70 are showed and color-coded according to the BED domain they belong to. The Zn^2+^ bound to the BED domain is pictured as a red sphere. E) Top: Curvature of LE11-27 DNA in the crystal structure. The color code for the DNA is the same as in A). The red line corresponds to the axis of the double helix obtained with Curves+ (Lavery et al. 2009). For clarity, only the residues 47-78 of the BED domain are shown. BED1 (dark grey) is at the front and BED2 (light grey) is in the back. Bottom: Plots of the depth of the grooves of LE11-27 in B-form (black) and bound to BED (blue) at the top strand bases A15 to T23. F) Mapping of the STRs 5’-AAGT-3’ (black arrowheads) and 5’-AAGC-3’ (red arrowheads) in the *Hermes* transposon Left End and Right End (LE and RE, respectively) as putative BED binding sites. The TIRs are pictured as blue arrows at the start of each end. The LE-STR1 and LE-STR2 are circled in red.

### The N-terminal domain of Hermes folds in a CCHC zinc-finger BED motif that specifically interacts with a subterminal repeat

Crystals were obtained of the BED/LE11-27 complex that were ultimately optimized to diffract to 2.5 Å, and the structure was solved by zinc multiwavelength anomalous diffraction (Zn-MAD). Iterative model building and refinement led to the atomic model presented in Fig. 2B. The crystal data, data collection and refinement statistics are reported in Table S1. The complex crystallized in space group P6_5_22 with a crystallographic twofold axis perpendicular to the oligonucleotide. Thus, despite the imperfect palindrome used in crystallization, the asymmetric unit contained one BED domain and one DNA strand, however the strand’s electron density was the superposition of the two not perfectly symmetrical DNA strands. This was likely due to the fact that the crystal lattice was held together by interactions between the blunt-ended DNA molecules packed end-to-end along the axis of the double helix. This resulted in a stochastic distribution of head-to-head and head-to-tail orientations that did not follow the lattice periodicity.

Each BED domain contains at its C-terminal a CCHC zinc-finger involving residues C51, C54, H71 and C73 that coordinate a bound Zn^2+^ ion. The two BED domains bound to the imperfect palindrome do not contact each other, but each C-terminal α-helix between T65 and R70 (α3; see Fig. 2B) is deeply inserted into the major groove. The summary of the protein-DNA interaction is presented in Fig. 2C. The two R70 residues in the assembly interact with the three bp at positions 17 to 19. The central three bp (positions 18 to 20) are not identical in the two orientations of the oligonucleotide in the crystals as bp 18 and bp 20 are either a C:G or an A:T pair, while bp 19 is a C:G but the bases flip positions between the two orientations. To accommodate for these variances, R70 and N67 are in two alternate conformations, consistent with the differences in H-bond patterns (Fig. 2D). These, together with S66, Q64 and the main chain carbonyl of R63 and T65 at the apical loop (between β2 and α3) of the zinc-finger, form all the base-specific interactions. In addition, there are several non-specific contacts between protein side chains of Lys, Arg, Ser, Thr and Trp residues and the DNA phosphate backbone. In one half of the palindrome, N67 interacts with the Hoogsteen face of A18, while R70 interacts with the stacked G17 and T18 on the complementary strand (Fig. 2D, BED1). In the other half (BED2), A18 is replaced by G20, which is less competent to interact with N67. We observe that the side chain of N67 is then flipped due to a steric conflict with R70, moving into its space to interact with G19 rather than G21. As a result, N67 is completely displaced, no longer facing the DNA but instead interacting with K59 and G19 phosphate (Fig. 2D, BED2). Thus, two Hermes BED domains can bind the two different DNA sequences of the imperfect palindrome due to the ability of the N67/R70 pair to assume alternate conformations.

Analysis of the DNA parameters indicates several distortions compared to B-form. The LE11-27 DNA experiences a slight bending of the double helix that is reflected in the positive roll and negative slide of base pair dyads (Fig. S2A and S2B). The bending is directed towards the edges of the major groove and the C-termini of the BED domains (Fig. 2E). Its minor groove shows widening over the base pairs flanking the central base pair (i.e., A16, G17, T18 and G20, C21, T22, respectively) of ~2 Å compared to B-form, whereas its major groove is narrowed around the central base pair before widening ~2 Å at the termini (Fig. S2C). On the other hand, the major groove is deeper by ~4 Å over the region A16-T22, while the minor groove is shallower over this region (Fig. 2E). This suggests that BED binding to the imperfect palindrome may be allosteric, in which the insertion of one α-helix into the major groove and the resulting DNA distortion makes the binding of the second BED easier.

### The mapping of the STRs

The interactions observed in the crystal structure of the BED/LE11-27 complex suggests that the BED binding site is 5’-AAG(T/C)-3’. There are 17 and 14 occurrences of this motif in the LE and RE of *Hermes* respectively, concentrated in particular towards the tips of the transposon, with five copies in the first 55 base pairs of both ends (Fig. 2F and Fig. S3). We designate these putative BED binding sites as subterminal repeats (STRs), and number them from “STR1” onward moving in from the LE transposon tip but from STR0 on the RE as the first AAGT motif is within the TIR. With the exception of LE-STR1/RE-STR1 and the pair LE-STR3/RE-STR2, the rest of the STRs of the two ends do not align with each other: their arrangement is asymmetric at the two ends. Interestingly, on the LE, except for STR1, STR2 to STR7 are all oriented antisense, while RE-STR1 to RE-STR5 are all oriented in the sense direction. Beyond LE-STR1/LE-STR2, only one other pair of STRs is organized as a palindrome, LE-STR14/LE-STR15, far interior in the LE beginning at bp 276.

### Reconstitution of the Hermes transpososome in vitro

As our attempts to crystallize either the full-length Hermes transposase or its complexes with transposon ends had failed, we turned to cryo-EM to determine the architecture of the transpososome, the assembly containing two bound transposon ends. Although the LE and RE TIRs only differ from each other by two base pairs, which the structures to date indicate are not involved with protein interactions, the transposase nevertheless has a much higher affinity for the LE compared to the RE (Hickman et al. 2014; Zhou et al. 2004). Therefore, to reconstitute the transpososome *in vitro*, we evaluated the interaction of various LE-DNA oligonucleotides complexed with the purified transposase to identify the best substrate for structural studies. We tested a range of oligonucleotides that extended the LE-TIR into the transposon such as LE-TIR+13 and LE-TIR+30 as well as those that were nicked and gapped at the non-transferred strand at cleavage site but also extended into the flanking DNA, i.e., 8+LE-TIR+7 and 8+LE-TIR+30 (Fig. 3A). Samples were prepared by mixing purified Hermes transposase with sub-stoichiometric or stoichiometric amounts of DNA and dialyzed to lower salt concentration prior to analysis by SEC.

**Figure 3:**
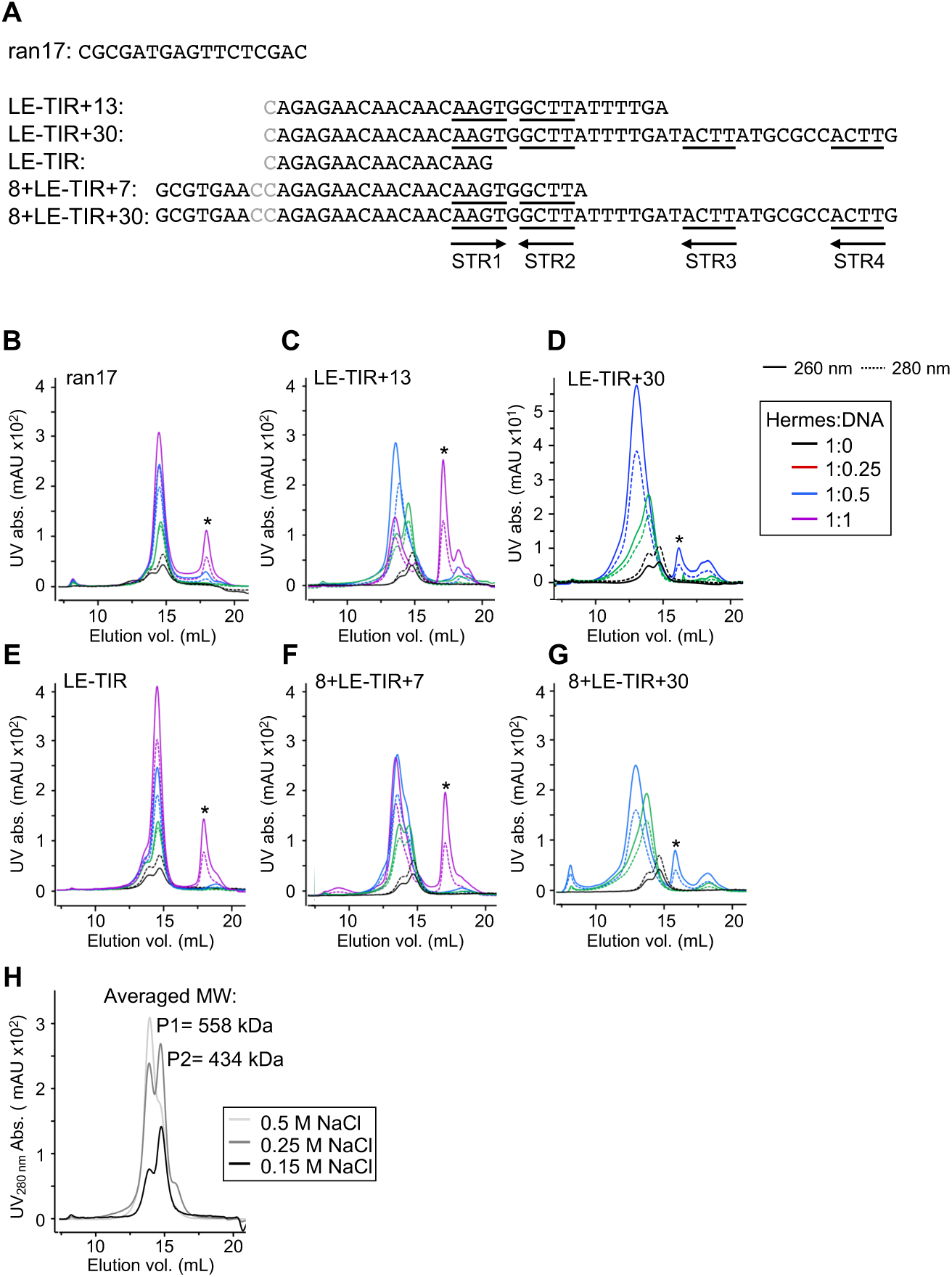
Analysis of the Hermes transposase in complex with various DNAs mimicking the Left End (LE) in its cleaved or nicked state by size exclusion chromatography (SEC). A) The sequence of the top strand of the double-stranded DNAs used in B). The bases in grey were absent on the top strand, but their complementary bases were present on the bottom strand. The subterminal repeats STR1 to STR4 are underlined and their orientation is indicated by arrows. B)-G) SEC chromatograms of the Hermes/DNA samples (buffer condition: 25 mM HEPES.Na pH 7.5, 150 mM NaCl). The Hermes-to-DNA ratios are shown in the insert. The free DNA elution peaks are marked by a star (*). H) SEC chromatogram of purified Hermes (in 0.75 M NaCl) dialyzed against three different elution buffers containing 0.5 M (light grey), 0.25 M (grey) and 0.15 M (black) NaCl. The apparent molecular weights (MW) derived from standard curve for two Hermes oligomeric populations, P1 and P2, are indicated.

The SEC elution profiles corresponding to titration of Hermes with the various LE-DNAs are presented in Fig. 3B-3G. The results were consistent with the crystal structure of the Δ-BED-transpososome (Hickman et al. 2014), in that the LE-TIR saturated the transposase with close to one DNA bound per protomer (Fig. 3E). However, for longer DNAs, the stoichiometry changed, with a maximum of 0.5 DNA bound per Hermes protomer (Fig. 3C, 3D, 3F and 3G). LE-TIR+13, LE-TIR+30 and 8+LE-TIR+30 led to precipitation of the complex when added to the protein in a stoichiometric 1:1 mix (Fig. 3C, 3D and 3G), in contrast to 8+LE-TIR+7 whose complex remained stable but did not bind more DNA in going from a 1:0.5 to 1:1 mix (Fig. 3F).

We also noticed that the peaks corresponding to the protein-DNA complexes were not monodisperse. This was also evident during the purification of the transposase, where we similarly observed heterogeneity and unusual behavior when the samples were analyzed by SEC. As shown in Fig. 3H, the purified transposase itself eluted as two populations, P1 and P2, where the relative ratio between the two elution peaks varied as a function of the salt concentration in the buffer, with the higher concentration favoring P1 and lower concentrations promoting P2. Through a combination of SEC and mass photometry (MP) analysis (Fig. 3H with Fig. S4), we established that the Hermes transposase exists in solution as two oligomeric states composed of six or eight monomers.

We used mass photometry (Fig. S5) to clarify the composition of the DNA-bound complexes observed in the chromatograms shown in Fig. 3B. These results were consistent with the SEC data showing that the Hermes transposase assemblies bound fewer molecules as the DNA was lengthened. For example, the MP experiments revealed that LE-TIR was mainly bound to the hexameric form of Hermes and that approximately five DNAs were bound. The longer oligonucleotides also generated more heterogeneous samples with the appearance of several masses under 400 kDa and broad distributions that led to poor fitting (standard error >30 kDa). Nevertheless, under our experimental condition, the Hermes hexamer could bind up to two of either LE-TIR+13 and 8+LE-TIR+30 DNAs; the Hermes octamer bound up to four LE-TIR+13, and three LE-TIR+30 and 8+LE-TIR+30 DNAs.

### Three LE-STRs bind to BED domains within the transpososome

On the basis of the SEC and MP experiments, for cryo-EM we selected the 8+LE-TIR+30 oligonucleotide as it contained both flanking region and the longest transposon DNA. Furthermore, it formed both hexameric and octameric complexes. As we assumed that the biologically relevant assembly is that in which one transposon DNA with its two ends is bound, we minimized the loading of the transposase and used a ratio of 1:0.25 protein-to-DNA. The resulting complex was purified by SEC and the fraction with the highest absorbance was used for structure determination.

As shown in Fig. 4A, the 2D classes of the top view of the complex resembled an octamer as observed in the Δ-BED-transpososome crystal structure (Hickman et al. 2014) with resolved features for two transposase dimers facing each other and linked by two disordered equatorial dimers to form a closed ring. The top views did not give clear indication for the presence of DNA, and were qualitatively very similar to the negative stained EM images obtained earlier on Hermes octamers without DNA (Hickman et al. 2014). However, in the side views, bound DNA was clearly visible. The particles were further cleaned by 3D classification and the best two classes (4 and 6) were refined and shown in Fig. 4B. The class 4 resembled an octameric Hermes that appeared to have lost one of its equatorial dimers, perhaps after binding DNA. The other classes were composed of closed octameric rings and as in the top view 2D classes, some of the equatorial transposase dimers were poorly resolved and not fully defined by the volume maps. Importantly, all the classes that we could refine presented two DNAs that bridged two Hermes dimers, the top “A” dimer and the bottom “C” dimer.

**Figure 4:**
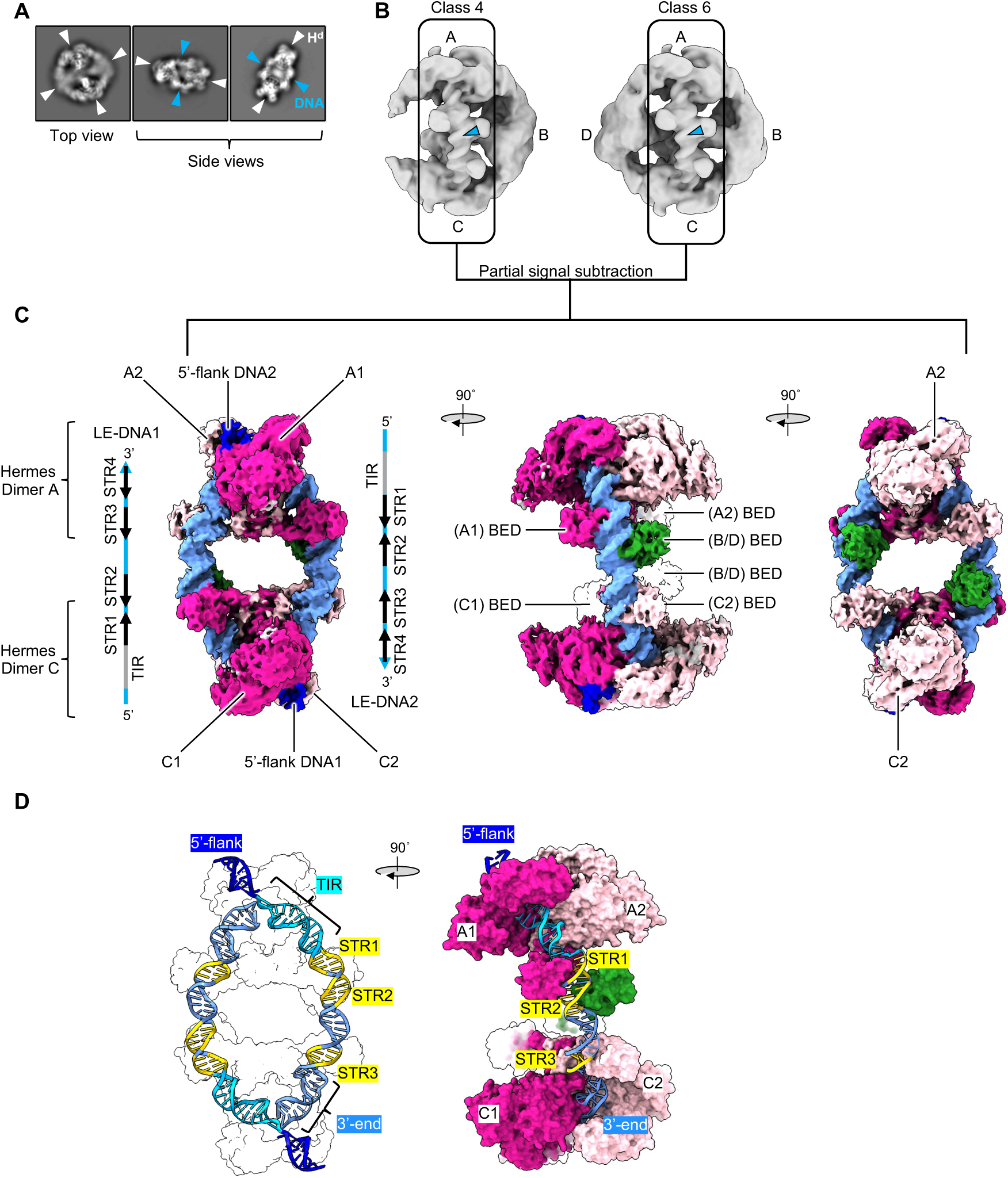
The cryo-EM structure of the Left End *Hermes* transpososome. A) Example of selected 2D classes of the transpososome from the data processing in RELION (Scheres 2012a, 2012b; Zivanov et al. 2018). The transposase dimers (H^d^) are marked with white arrowheads and the DNAs are indicated with blue arrowheads. The box size is 280 Å. B) Selected 3D classes 4 and 6 from RELION processing (top views). The transposase dimers are indicated as A, B, C and D, and the DNAs visible on the displayed faces are marked with blue arrowheads. The boxes enclose the regions (Hermes dimers A and C as well as the DNAs and the DNA-bound BED domains) that were best resolved. The particle images were partially signal subtracted to remove the poorly resolved Hermes dimers B and D, focusing the reconstruction on the core of the transpososome. C) The gold-standard refined map of the core of the transpososome at 4.64 Å resolution. The full Hermes protomers are colored in pink. The light pink protomers A2 and C2 interact with the 3’-end of the DNAs, and the bright pink protomers A1 and C1 interact with the 5’-end of the DNAs. The DNAs are in blue, with their 5’-flank in dark blue. BED domains belonging to the dimers B or D are in green. The feature organization of the LE-DNAs (TIR and STRs) is shown schematically on each side of the map. D) The atomic model of the core of the LE/LE *Hermes* transpososome (PDB: 8EDG). Left: the antiparallel orientation of the LE-DNAs inside the complex (only the silhouette of the atomic surface of the Hermes dimers is shown). The features of the DNAs are highlighted, but only the STRs interacting with the BED domains are reported. Right: the atomic surface of the Hermes dimers (same color coding as in C) is displayed while the LE-DNA2 is shown in cartoon mode to emphasize the extended protein/DNA interaction inside the transpososome.

To optimize the alignment of the most resolved regions of the transpososome, we performed partial signal subtraction on the best 3D classes using a mask that excluded the poorly resolved region of the equatorial Hermes dimers (units B and D in Fig. 4B). As a result, the final map (Fig. 4C) corresponds to the core of the LE-LE-transpososome two Hermes dimers A and C (with each monomer A1/A2 and C1/C2 in light and bright pink) are bridged by two DNA oligonucleotides (in blue). Also shown in Fig. 4C are two DNA-bound BED domains (in green) that are contributed by the equatorial dimers which were partially masked-out during the processing. Overall, three BED domains bind to each DNA. It was possible to unambiguously assign the orientation of the DNA molecules (hereafter “5’” and “3’” always refer to the top strand) since the 5’-flanking region as well as the two-nucleotide gap on the top strand of the 8+LE-TIR+30 DNA were clearly visible. The two LE DNAs are antiparallel, with each TIR interacting with a different Hermes dimer next to the 3’-end of the other oligonucleotide (Fig. 4C). The configuration of the transpososome with two antiparallel DNAs was the only configuration we saw. Based on these observations, we rigid-body fitted into the potential map several copies of the crystal structure of the N-truncated Hermes dimer bound to two 8+LE-TIR oligonucleotides (PDB: 6DX0) (Hickman et al. 2018), and that of the BED/LE11-27 complex (Fig. 4D).

The 8+TIR-LE+30 oligonucleotide includes the TIR and STR1 through STR4. STR1 is clearly bound by the BED domain that belongs to the same Hermes protomer (A1 or C1) that also bind to the TIR (bright pink) as indicated by distinct linker density between the BED domain and the dimerization domain (DD in Fig. 1C). The identity of the equatorial Hermes protomers that supplied their BED domain to each STR2 was unclear. However, before we masked out the equatorial dimers, refinement of the initial 3D classes of Fig. 4B at 6.25 Å resolution showed weak densities that linked these BED domains to the equatorial dimers (Fig. S7). The connectivity between the BED domains and the equatorial dimers varied depending on which 3D class was used, suggesting that in solution different species with different connectivities may exist. The final map also showed weak density that linked the STR3-bound BED domains to the core of protomers A2 or C2 (light pink). STR4 was not bound by any BED domain; rather, the 3’-end of each oligonucleotide (bp 36-47) was positioned similarly as the TIR entering the catalytic domain of the A2 or C2 protomer. Of note, no protein-protein interaction between the BED domains or between the BED domains and the rest of their transposase was observed.

### The STRs of the RE do not interact with the BED domains inside the transpososome

As we were unable to generate interpretable cryo-EM data for a single Hermes assembly that bound one LE and RE transposon end at the same time, we determined the structure of the transpososome bound to two REs using the same approach as described for the LE-LE version. Despite extensive efforts to optimize the cryo-EM specimen preparation and data processing, the reconstruction only achieved 5.1 Å resolution. The 3D class that refined best clearly showed 5’-flanks synapsed in the top A1/A2 dimer revealing two parallel DNAs. Strikingly, in none of the 3D classes did we observe density corresponding to either free or bound BED domains (Fig. 5). Both 3’-ends of 8+RE-TIR+30 were inserted in the C1/C2 dimer in a TIR-like fashion, suggesting that this interaction was not a function of STR/BED interactions, but a possible consequence of the particular length of the DNA used.

**Figure 5:**
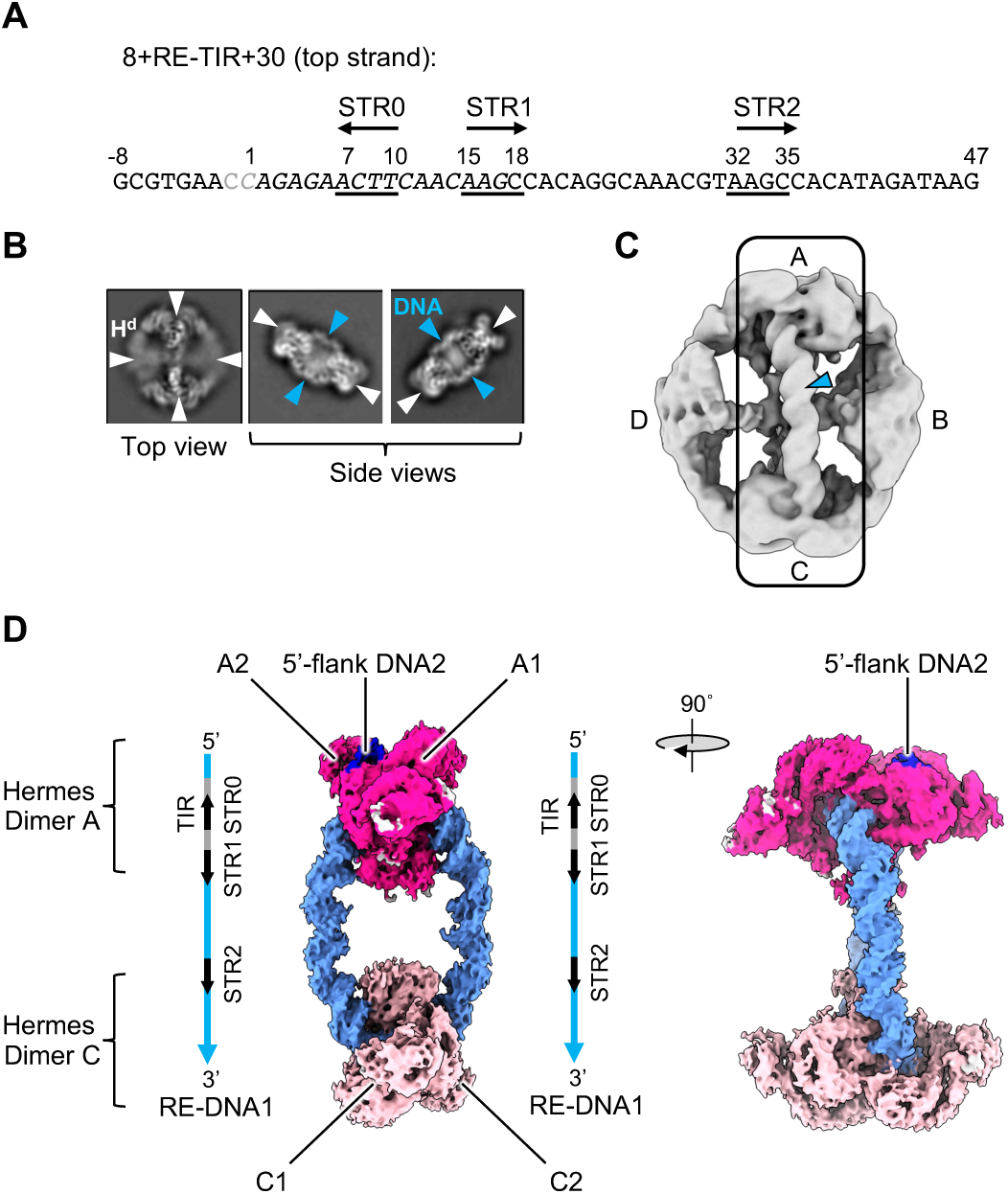
The cryo-EM structure of the Right End *Hermes* transpososome. A) Sequence of the top strand of the 8+TIR-RE+30 DNA (RE-DNA). The bases in gray were absent on the top strand but their complements were present in the bottom strand. The terminal inverted repeat (TIR) is in italic, the subterminal repeats STR0 to STR2 are underlined, and their orientation indicated by arrows. B) Example of selected 2D classes (box size is 250 Å) from the data processing in RELION (Scheres 2012a, 2012b; Zivanov et al. 2018). The transposase dimers (H^d^) are marked with white arrowheads and the RE-DNAs are indicated with blue arrowheads. C) The best 3D class of the transpososome (top view). The transposase dimers are marked A-D, and the RE-DNA visible on the displayed face is marked with a blue arrowhead. The particle images were partially signal subtracted to remove the poorly resolved Hermes dimers B and D, focusing the reconstruction on the core of the transpososome (contained inside the black box). D) The gold-standard refined map of the core of the RE/RE transpososome at 5.1 Å resolution. The Hermes dimers A1/A2 and C1/C2 are colored in bright and light pink, respectively. The RE-DNAs are in blue, with their 5’-flank in dark blue, signaling their parallel orientation. The protomers A1 and A2 interact with the 5’-end of the DNAs, and the protomers C1 and C2 interact with the 3’-end of the DNAs. No clear density for DNA-bound or free BED domains is present. The features of the RE-DNAs (TIR and STRs) are pictured on each side of the map.

### The minimal BED binding site is the AAGT motif, but it must be followed by an AT-rich region

It was very surprising that in the RE-RE complex, no BED domain was bound to RE-STR1 as we had anticipated it would be bound as seen for LE-STR1. This forced us to reevaluate the notion that the AAG(T/C) motif indeed represents the BED domain recognition site. We first asked whether RE11-27 that contains RE-STR1 (5’-AAGC) binds the BED domain (Fig. S9). SEC analysis showed that RE11-27 DNA binds similarly as the randomized control, ran17, and did not form a stable complex. Likewise, a mutant RE11-27 (RE11-27T) with an AAGT motif replacing AAGC did not form a stable complex either. However, on the LE, LE11-27mut where STR2 (5’-AAGC) was replaced by 5’-AAAA formed a stable 1:1 protein-to-DNA complex. These results suggested that an isolated AAGT motif was necessary but not sufficient for BED binding. We noted that the AAGT motif in LE11-27mut was flanked by two AT-rich regions, whereas RE11-27T presented the AT-rich region only at its 5’-end. We then tested if a BED domain interacts with an isolated AAGT motif only if it is followed by an AT-rich region (Fig. S9, LE11-27mut-5’G vs. LE11-27mut-5’3’G), and we found this was indeed the case. We verified whether it was also the case for an isolated AAGC motif as the RE-STR1 did not show binding. The SEC data obtained with LE11-27mutC showed only weak binding comparable to ran17, suggesting that an isolated AAGC motif was not sufficient. Collectively, these experiments revealed that the AAGT motif is the effective BED binding site, but it must be accompanied by an AT-rich region to achieve high affinity DNA binding.

### The interaction of the BED domains with the quasi-palindromic LE-STR1-STR2 is cooperative

The characterization of the BED binding site suggested that the AAGC motif might need the presence of the AT-rich AAGT motif adjacent to it for optimal binding. Furthermore, the crystal structure suggested the possibility of cooperative binding between the two BED domains as two α3 helices from two protomers are inserted into a deepen major groove adjacent to each other. To test this hypothesis, we used electrophoretic mobility shift assays (EMSA) with fluorescently labelled LE11-27 and LE11-27mut DNAs and the BED domain. The resulting polyacrylamide gels are shown in Fig. 6B. As expected, the DNAs generated two different delayed bands resulting from the interaction of two and one BED with LE11-27 and LE11-27mut, respectively. No band corresponding to a single binding event, even in the early stage of the titration, was detected for LE11-27. Therefore, only simultaneous double interaction events were observed consistent with cooperative binding. The EMSA data for LE11-27 were converted to a Hill plot (Fig. 6C), and the value of the slope coefficient (n_H_ = 1.7) confirms the positive cooperative binding of BED. As the crystal structure of the BED/DNA complex showed no evidence of protein-protein contacts, it appears therefore that cooperativity is the result of DNA deformation.

**Figure 6:**
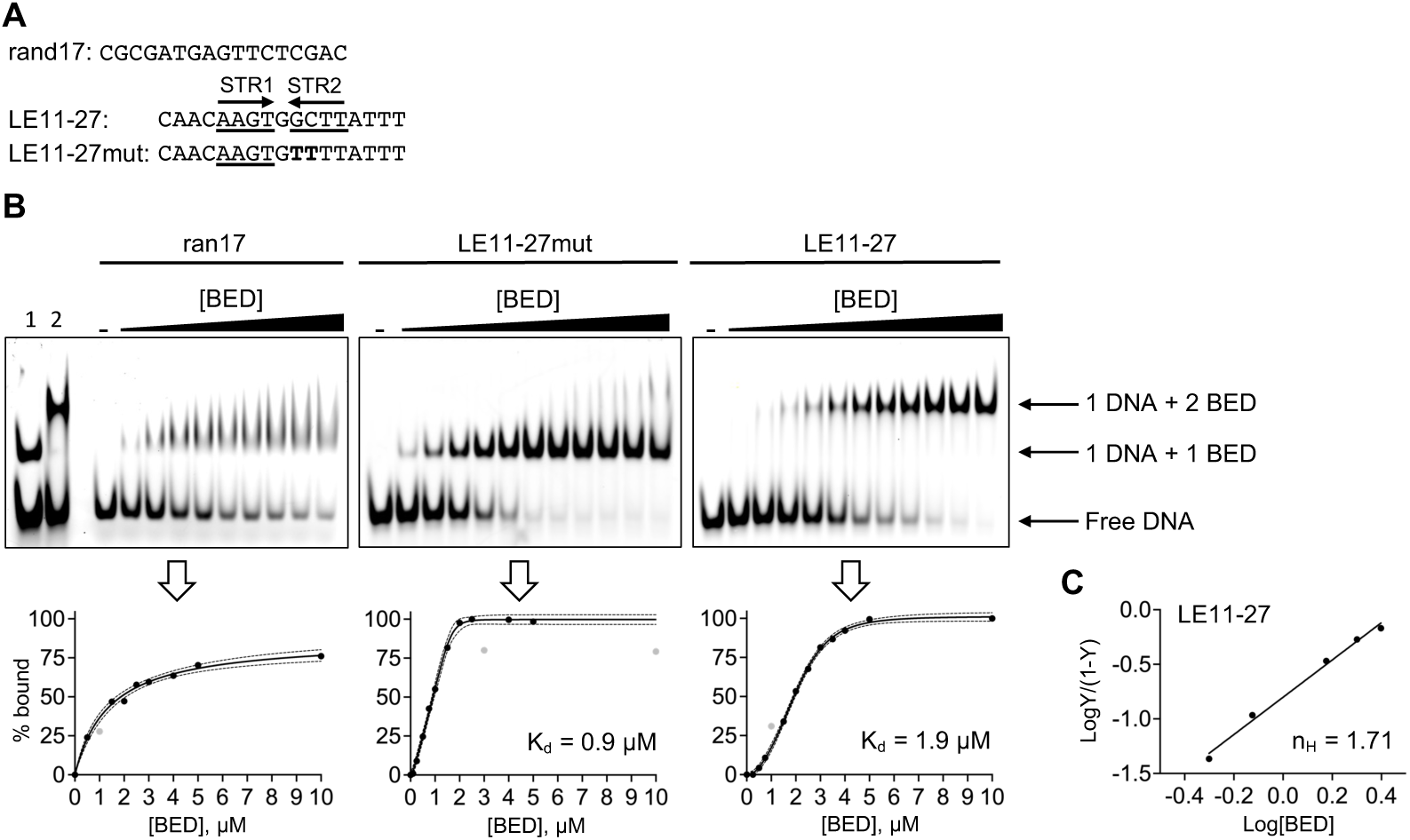
The cooperative interaction of the BED domain with the *Hermes* transposon Left End (LE) quasi palindromic STR1-STR2 motif revealed by electrophoretic mobility shift assay (EMSA). A) The sequence of the top-strand of the DNAs ran17, LE11-27mut and LE11-27 double-stranded DNAs are presented at the top of the EMSA polyacrylamide gels (15% polyacrylamide, 0.5 X TBE). The subterminal repeats STR1 and STR2 are underlined, and their orientation is indicated with arrows. The mutated bases in LE11-27mut are in bold. Ran17 is the unspecific interaction control. B) EMSA polyacrylamide gels. The DNAs (1 µM) were titrated with increasing amount of BED protein from 0 (-) to 10 µM. The lanes 1 and 2 in the ran17 EMSA correspond to a mix of BED with LE11-27mut and LE11-27 as markers, respectively. Under each gel, the percentage of bound DNA derived from the band intensity is plotted against the concentration of BED domain. The data points were fitted, and the dash lines describe the 95 % confidence interval of the fitting. The data points in grey were treated as outliers and excluded from the fitting. The dissociation constants (K_d_) derived from the fitting are reported for LE11-27mut and LE11-27. C) Hill plot for the binding of LE11-27, Y/(1-Y) being the proportion of bound DNA. The Hill coefficient (n_H_) with a value greater than 1 describes the positive cooperative binding of BED.

### The formation of the synaptic complex does not only rely on the asymmetry of the first STRs in cells

The cryo-EM reconstructions indicated that the LE/LE pair, in contrast with the RE/RE pair, is in antiparallel orientation. We suspected that the BED/STR interactions within the LE/LE pair were responsible for the antiparallel orientation. Previously, we showed that a modified *Hermes* transposon with symmetrized LE/LE ends were inactive in insect cells (Hickman et al. 2014). We asked if we could rescue the transposition of a LE/LE transposon carrying a puromycin marker, by mutating the first three STRs of one end to make it resemble the RE, contained on a LE-mut donor plasmid, p-donor (Fig. 7A-7B). The transposase was expressed from a helper plasmid, p-helper. Both plasmids were transfected into HEK293 cells. After several days under puromycin selection colonies were counted as a read-out (Fig. 7C). As *Hermes* transposition in HEK293 cells has not been previously reported, we first established it was active on its wild-type ends. As a comparison, *Hermes* showed a higher transposition rate than that of wild-type *piggyBac* under identical transfection conditions. (*Hermes* was also reported as highly active in *S. pombe* so that the T317A mutation had to be introduced to tame it (Evertts et al. 2007)). Symmetrized LE/LE p-donor led to a drastic loss of mobilization in *Drosophila* S2 DEV8 cells (Hickman et al. 2014) and, similarly, in HEK293 cells, LE/LE p-donor retained little mobilization, while the RE/RE transposon showed no activity. Finally, the LE/LEmut p-donor showed the same activity as the LE/LE p-donor. Thus, restoring the STR asymmetry in the first 50 bp in one of the LEs was not sufficient to rescue the mobilization of the LE/LE transposon. Similarly to previous experiments in insect cells, in cell transposition activity requires substantially longer transposon ends that is not feasible to study with high resolution experimental tools. However, it appears clear that the asymmetry of the transposon ends is an important requirement for activity.

**Figure 7:**
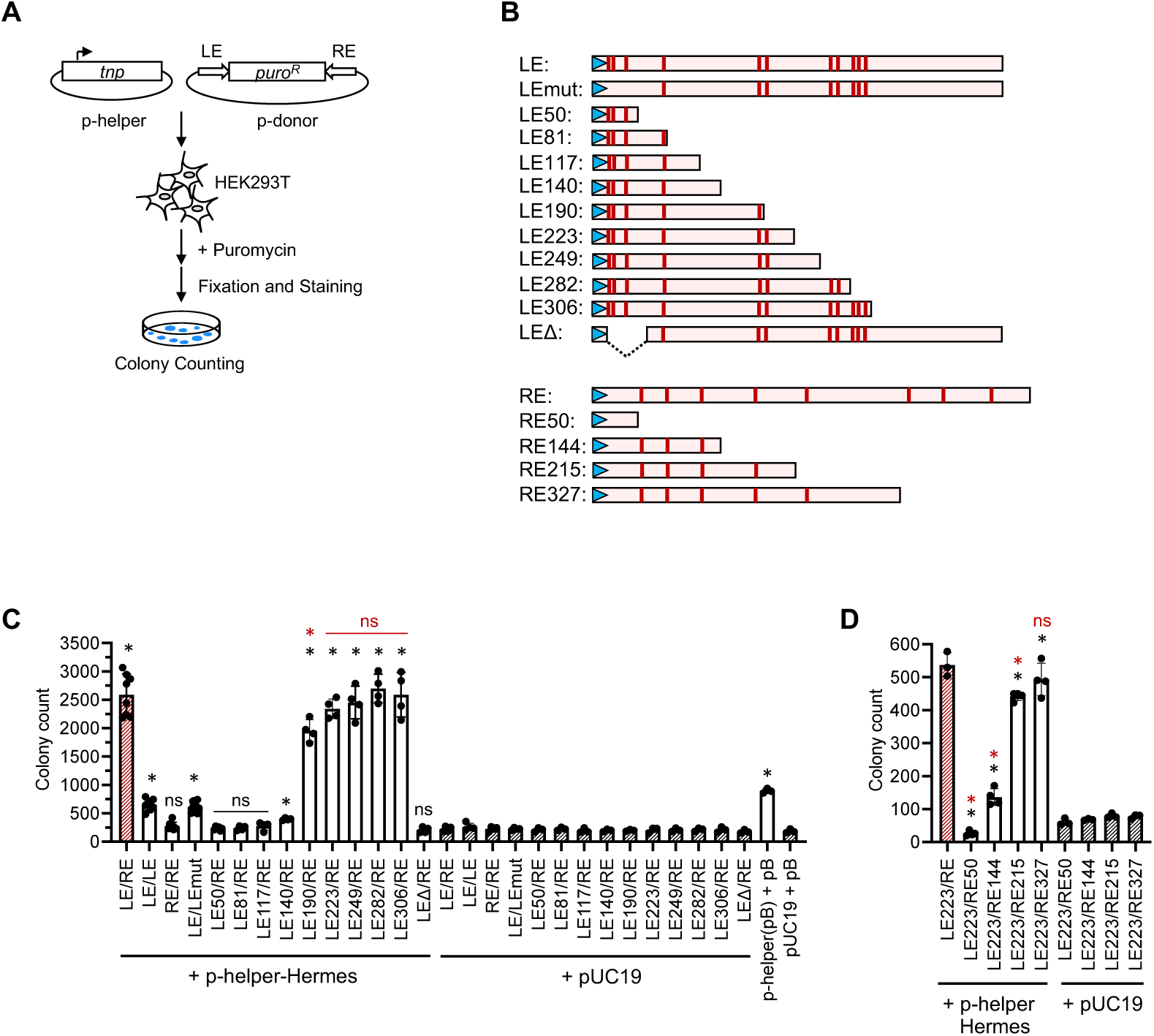
*Hermes* transposition assay in HEK293T cells. A) Schematic of the experimental procedure. The plasmid p-helper encoding the Hermes transposase (tnp) and the p-donor containing the puromycin resistance gene puro^R^ (1.5 kb) flanked by the *Hermes* transposon Left End (LE) and Right End (RE) are transfected into HEK293T cells. The cells that experience a chromosomal integration of the LE-puro^R^-RE transposon are selected against puromycin and the colonies counted. B) Schematic of the *Hermes* LE, RE and their variants (truncation or mutations) used in the assay. The Terminal Inverted Repeats (TIRs) are represented by blue arrowheads and the subterminal repeats putative Hermes BED domain binding sites are depicted as red bars. C)-D) Histograms of the colony count for each p-donor. The combination of the transposon ends is reported on the x-axis for each p-donor. Each black circle represents one independent data point, with 4 < n < 7 for the assay (white columns) and n ≥ 3 for the controls (red stripped and black stripped columns). The mean values are plotted as columns and the standard deviations are represented with bars. The statistical t-test was applied to determine whether the experimental data were statistically different from their pUC19 controls (black star and black ns) or from the LE/RE experiment (red star and red ns). The *piggyBac* transposition system (pB) was used as a general control of the experimental design.

### The minimal end requirement for Hermes transposition in HEK293T cells

The transposition assay showed that the divergence of sequence between the ends must span beyond the first 50 bp to have a mobile system. Hence, we sought to establish how much of the LE is needed to observe transposition while keeping the RE unchanged. We tested nine different LE/RE p-donors wherein the LE was gradually truncated (LE50 to LE306) as well as one in which the region spanning over the first three STRs was deleted (LEΔ) (Fig. 7A), the results are presented in Fig. 7C. The respective lack of significant integration of the transposons LE50/RE and LEΔ/RE confirmed that neither the 50 bp downstream the LE-TIR nor the rest of the LE are sufficient for mobilization in cells when combined with the full RE.

The truncated LE constructs showed that mobility equivalent to the full-length LE was retained up to 223 bp. A slight decline in integration started at 190 bp to finally reach a drastic downturn at 140 bp and shorter. The loss of transposition activity was steady, allowing us to affirm that the critical truncation point on the LE lies between 140 and 190 bp when associated with the full-length RE. Experiments with partially scrambled LE sequences suggest that there is no influence on transposition by not keeping constant the distance between the LE-TIR and the CMV enhancer (Fig. S10). Similarly, we also asked what the minimal end requirement was for the RE when combined with LE223 (Fig. 7D). We observed that the RE truncated after 327 bp resulted in similar integration as the full-length RE. The transposition activity is slightly reduced with 215 bp of RE, while it is severely impaired when only 144 bp of RE were retained. We conclude that the *Hermes* transposon needs at least ~220 terminal bp of both its LE and RE to be effectively mobilized in cells.

## Discussion

The cut-and-paste *Hermes* transposon stands out among the transposases that have been biochemically or structurally characterized so far. While most DNA transposases function as dimers or tetramers, Hermes is unique that it forms closed octameric rings as tetramer of dimers. While a mutant version of Hermes, incapable of forming octameric rings, that forms only dimers is hyperactive for transposition *in vitro* at low salt conditions, it was inactive at physiological ionic strengths or in cells (Hickman et al. 2014). We sought to establish a mechanistic explanation of the higher order organization necessary for in cell activity.

The cryo-EM structures of the cores of the LE-LE and RE-RE transpososomes determined here provide a new framework for understanding the role of Hermes’ unique ring-shaped organization. In the LE-LE complex, although the two ends were antiparallel (an arrangement that clearly cannot support transposition), each of the first three STRs within the first 30 bp of the LE interacted with a BED domain originating from three different Hermes dimers. This includes the high affinity LE-STR1-STR2 site that is present only on the LE of *Hermes.* The RE adopted a similar but parallel positioning within the transpososome, but it did not interact with BED domains Fig. 8A. Taken collectively, we propose that transposition activity in cells relies on a transpososome assembly that can supply sufficient number of BED domains and in the correct configuration to first recognize and tightly bind to its transposon LE. Seemingly, a ring-shaped octamer that assembles a stable closed form oligomer has evolved to serve this purpose. Furthermore, the binding of the first end, as it interacts with three dimers out of four, might shape the three-dimensional organization of the Hermes assembly, that might not be able to accommodate three STR/BED interactions with a second end parallel to the other. Thus, the antiparallel LE binding might be the most energetically stable solution for the Hermes assembly. However, such arrangement does not support transposition activity. We propose that the cooperative binding of two BED domains to the LE-STR1-STR2 palindrome is the key to the recognition of the LE and this is the first binding event. The cryo-EM structure of the core of the RE-RE transpososome showed that the RE does not interact with any BED domain, suggesting that RE binding solely relies on the interaction of its TIR. Assisting the process is the fact that once the LE is bound, the RE is only a few kb away at the other end of the transposon (Fig. 8B).

**Figure 8:**
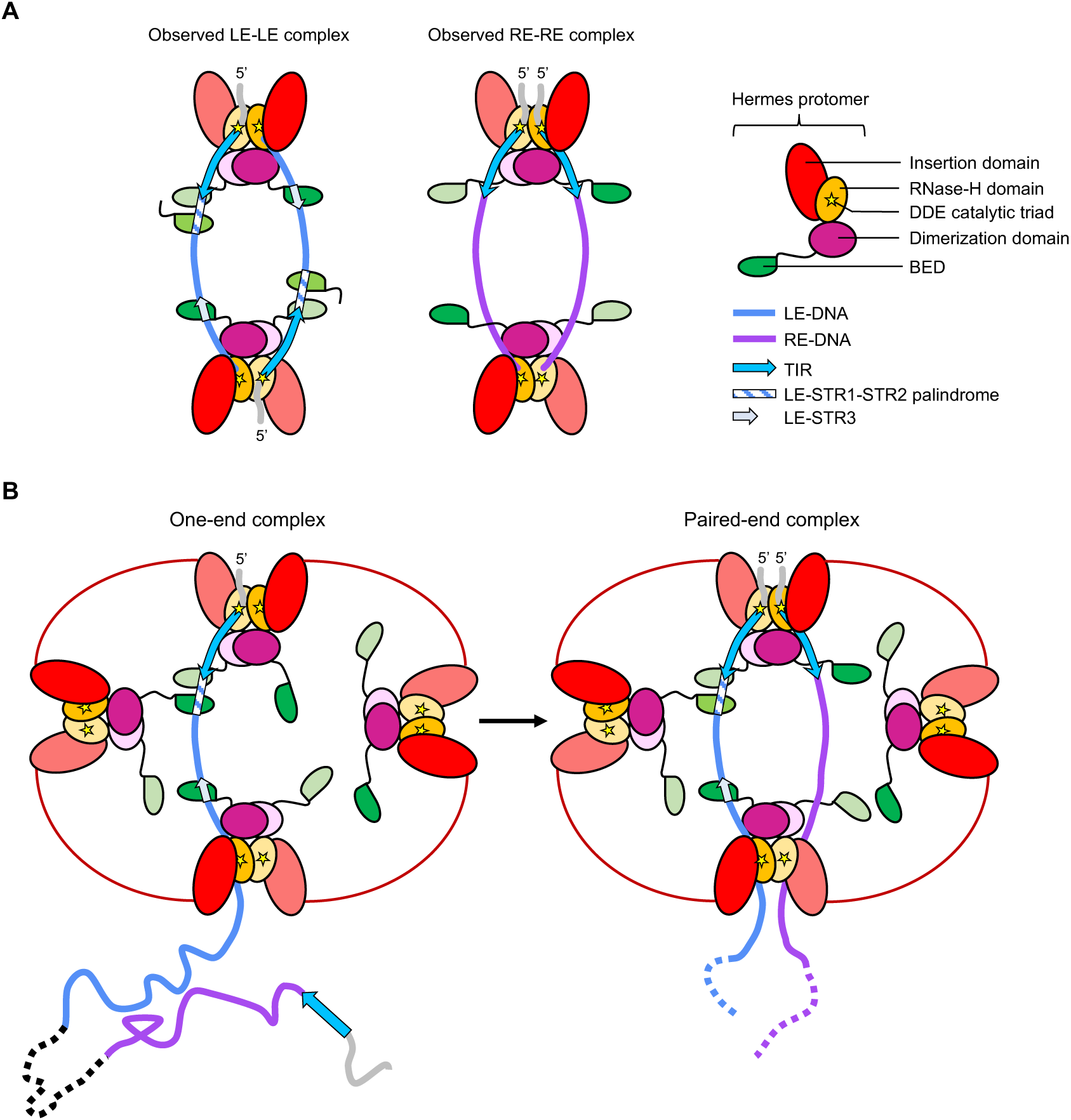
The model of the formation of the *Hermes* transpososome in cells. A) Schematic of the organization of the core of the *Hermes* LE/LE (left) and RE/RE (right) transpososomes observed by cryo-EM. In the LE/LE transpososome, the LE-TIR interacts with the catalytic center of one Hermes protomer, the LE-STR1-STR2 palindrome and the LE-STR3 interact with three BED domains from three Hermes protomers belonging to three dimers, and the LE-DNA 3’-end interacts with the opposite Hermes dimer. The LE-DNAs interact with the transposase assembly in antiparallel orientation. In the RE/RE transpososome, the RE-TIR interacts with the catalytic center of one Hermes protomer, and the RE-DNA 3’-end interacts with the Hermes dimer. The RE-DNA does not interact with BED domains. The RE-DNAs interact with the transposase assembly in parallel orientation, with both RE-TIRs present in the same Hermes dimer. B) Proposed model of the formation of the *Hermes* transpososome in cells. The LE is first recognized by the transposase assembly, because of its higher affinity conferred by its quasi-palindromic STR1-STR2 motif (one-end complex). Once the LE is bound, the RE is only a few kilobase away at the other end of the transposon and can then be recognized solely based on the sequence its TIR. Both TIRs are synapsed inside the same Hermes dimer (paired-end complex) and the transpososome is in the proper configuration to perform its cut-and-paste transposition chemistry.

The LE-STR/BED interaction relies on a rigid pattern with the STR motifs precisely positioned relative to one another and to the TIR, compatible with the three-dimensional arrangement of the BED domains inside the transposase octamer. For example, the cooperative binding of two BED domains with the LE-STR1 and LE-STR2 is only possible due to their palindromic arrangement, and we have previously shown that even a single bp shift in the position of the LE-STR1 and LE-STR2 away from the LE-TIR severely impairs the transposon cleavage activity (Hickman et al. 2014). These observations suggest that the exact phasing of the LE-STRs is crucial. Certainly, the closed architecture of a ring-shaped assembly is more rigid when compared to the alternative of linear arrangement, and we suggest that this is an important aspect of supporting the proper spatial organization of the BED domains and their interactions with the LE-STRs.

Conceptually, the accumulation of zinc-finger (ZF) motifs for high-affinity binding is similar to that of transcription factors (TFs) that are composed of ZF arrays. For example, the essential insulator protein CTCF has eleven ZFs (Yin et al. 2017), and KRAB-ZF proteins feature twelve ZFs on average (Urrutia 2003). The ZFs of these polydactyl proteins recognize a wide variety of DNA motifs of typically three or four bases avoiding redundancy within the same protein. The organization in arrays enables the combination of ZFs to interact with longer target sequences and increases the affinity of the protein towards DNA sites. As the Hermes transposase did not evolve the luxury of multiple BED domains arranged as beads on a string as did ZF-TFs, it apparently solved its affinity problem by high order multimerization to make an array out of a single domain on the polypeptide. The use of a short recognition motif has a problem, though, as it is present through the genome. Thus, while the 5’-AAGT motif is the essence of the minimal BED binding site, high affinity interaction occurs only if a AT-rich region follows.

It was previously suggested that the *Hermes* STR was 5’-GTGGC as it was repeated several times close to the TIRs, and also found in the closely related *hobo* transposon (Hickman et al. 2014; Kim et al. 2011). Our binding data suggests that this should be reevaluated, as 5’-AAGT represents the BED binding site although its binding mode is complex as it also depends on the sequence following the tetranucleotide. The Hermes BED domain has some nonspecific DNA binding affinity but reaches high affinity specific binding only if the AAGT motif is followed by an AT-rich sequence. Under these constraints, *Hermes* has at most eight other putative BED binding sites on the LE after bp 50, and eight on the RE (Fig. S2B). The *Hermes* ~200 bp minimal ends requirement for in cell transposition strongly suggests that the sequence beyond the terminal 50 bp of both ends is involved in the formation of a competent transpososome. However, it remains unclear whether more STRs are involved in the assembly of the transposition complex. It is also possible that the chromatin architecture may play a role in the process. Interestingly, there are two putative BED binding sites between the LE bp 140 and 223 at the positions 183 and 190 (LE-STR5 and LE-STR6, respectively). Two BED binding sites were also predicted in the RE between the bp 144 and 327 at the positions 172 and 228 (RE-STR4 and RE-STR5, respectively). These motifs might be important for the formation of the transposition complex in cell. We note that the distance between the LE-STR3 and the LE-STR5 is 147 bp, which exactly corresponds to the length of DNA packed around a nucleosome. The presence of a nucleosome could induce the LE to fold back bringing the LE-STR5 and LE-STR6 close to the TIR-bound transpososome. The RE could similarly carry one or several nucleosomes whose DNA bending action may help in the synapse of the *Hermes* transposon ends. Similar observations have been made for the *piggyBac* transposon, for which there is a left-internal site present at one nucleosome length from the *piggyBac* LE-TIR (Morellet et al. 2018).

Our results have implications beyond *Hermes* to other members of the *hAT* superfamily. Kunze and coworkers have reported that the N-terminal region of the *hAT* Activator (Ac) transposase recognizes several STRs scattered in both ends of *Ac* in a cooperative manner (Becker and Kunze 1997; Feldmar and Kunze 1991; Kunze and Starlinger 1989). We suggest that it is the cooperative binding of Ac’s C_2_H_2_-BED domains that might be responsible for this. Most of the *hAT* transposons have sets of STRs close to the TIRs (Atkinson 2015). Interestingly, the *hAT Tol2* transposon presents five 5’-AAGT motifs in the first 50 bp of its LE and RE, and the LE somewhat resembles *Hermes* LE, featuring a perfect palindrome (5’-AAGTACTT) directly after the TIR followed by a third anti-sense STR 10 bp downstream (Koga et al. 1999). Furthermore, all the *hAT* transposases have a BED domain at their N-terminus (Atkinson 2015). However, to our knowledge Hermes is the only *hAT* transposase that was expressed and purified to obtain soluble transposase; therefore, biochemical and biophysical data involving the crucial question of multimerization by other *hAT* transposases are unfortunately not currently available. It has been suggested that Tol2, Tgf2 and the domesticated Kat1 can form oligomers (bigger than dimers) spontaneously or upon DNA binding (Chiruvella et al. 2016; Jiang et al. 2016; Shibano et al. 2007). Therefore, it is possible that the ability to deploy a multitude of BED domains and to cooperatively bind to STRs might be generalizable to other *hAT* transposons as well.

The *hAT* transposons have been co-opted several times as a source of coding sequence for the emergence of new genes and functions (Atkinson 2015). Even though domesticated proteins have evolved from intact transposases, it seems that their enzymatic activity is rarely retained contrary to their DNA binding capacity (Etchegaray et al. 2021; Sinzelle, Izsvák, and Ivics 2009). The *hAT* BED domain has been conserved in many cases and even sometimes replicated as in the vertebrate ZBED4 and ZBED6 proteins that have four and two BED domains, respectively (Hayward et al. 2013). The ZBED transcription factors are expressed in various vertebrate tissues and have been found to be involved in the regulation of many functions such as the expression of ribosomal protein genes (Yamashita et al. 2007), embryogenesis and carcinogenesis (Hayward et al. 2013), retinal morphogenesis (Mokhonov et al. 2012) or muscle development (Chen et al. 2009). Our structural results might help in better understand how these proteins function.

The simplicity of cut-and-paste transposon systems makes them appealing to re-purpose as genomic tools as transposons are naturally occurring genome editing systems. Importantly, cut-and-paste DNA transposition does not require extensive DNA repair and therefore can work in cell lines where repair mechanisms are not efficient. Currently, the two most widely used transposon systems to modify mammalian genomes are *piggyBac* (Kim and Pyykko 2011) and *Sleeping Beauty* (Kebriaei et al. 2017). However, as wild-type transposases typically have suboptimal activity since they are not under positive selection, this gives us the opportunity to generate hyperactive transposons. A spectacular example of this possibility is *Sleeping Beauty*, whose transposase was first reactivated and then at least two orders of magnitude of increased transposition activity was achieved using educated guesses from sequence alignment and randomized genetic screens (Ivics et al. 1997; Mátés et al. 2009). The *Hermes* transposon is mobile in a variety of organisms from yeasts (Gangadharan et al. 2010; Park, Evertts, and Levin 2009; Patterson et al. 2018) to various insects (Guimond et al. 2003; Pinkerton, O’Brochta, and Atkinson 1996; Sarkar, Coates, et al. 1997; Sarkar, Yardley, et al. 1997) and here we showed also in human tissue culture and at activity levels comparable to *piggyBac*. The understanding of the principles of transpososome assembly and organization at the three-dimensional level should open up possibilities for the rational redesign of Hermes to join the array of transposon-based tools available for the modification of mammalian genomes.

## Material and Methods

### Protein expression and purification

#### Isolated BED domain

The region coding for the N-terminal BED domain of Hermes (Uniprot Q25438) spanning from residues 1 to 78 (BED) was cloned into a pET15b expression vector in such a way that the recombinant protein was not tagged. The protein BED was expressed in Rosetta2(DE3) *E. coli* cells (Novagen) by growth at 37 °C in LB medium supplemented with 50 µg/mL carbenicillin until reaching OD_600nm_ ~0.6, followed by cooling at 16 °C and induction by addition of IPTG to the final concentration of 0.25 mM. After 16 hours, 1 L of culture was harvested by centrifugation and resuspended in 50 mM Tris-HCl pH 7.5, 100 mM NaCl, 0.3 mM TCEP, 2.5 mM MgCl_2_, DNaseI and protease inhibitors (Roche). The cells were disrupted by sonication and the soluble part of the lysate was purified on a 5 mL-HiTrap heparin column (GE Healthcare) at 4 °C. The column was equilibrated with 25 mM HEPES.Na pH 7.5, 100 mM NaCl, 0.3 mM TCEP and protease inhibitors. BED was eluted with a linear salt gradient (100 mM to 1 M NaCl). The fractions that contained BED were concentrated and further purified at 4 °C on a preparative Superdex 75 16/600 size exclusion column (GE Healthcare) equilibrated with 25 mM HEPES.Na pH 7.5, 500 mM NaCl and 0.3 mM TCEP (see chromatogram in Fig. S1A). The protein and its purity were checked at each step of the purification by SDS-PAGE electrophoresis (4-12% Bis-Tris NuPAGE, MES as running buffer, Invitrogen) stained with SimplyBlue SafeStain (Novex) (Fig. S1A).

#### Hermes transposase

The region coding for the full-length Hermes transposase (Uniprot Q25438, with the mutations Q2E and K128G) was cloned into pBAD/Myc-His (Invitrogen) in such a way that the recombinant protein was not tagged. Hermes was expressed in Top10 (Invitrogen) *E. coli* cells by growth at 37 °C in LB medium supplemented with 50 µg/mL carbenicillin until reaching OD_600nm_ ~0.6, then followed by cooling at 16 °C and induction by addition of arabinose to the final concentration of 0.01 %. After 16 hours, 4 L of culture were harvested by centrifugation and resuspended in 50 mM Tris-HCl pH 7.5, 500 mM NaCl, 0.3 mM TCEP, 40 µM MgCl_2_, DNaseI and protease inhibitors (Roche). The cells were disrupted by sonication and the soluble part of the lysate was purified on two 5 mL-HiTrap heparin columns (GE Healthcare) at 4 °C. The column was equilibrated with 25 mM HEPES.Na pH 7.5, 100 mM NaCl, 0.3 mM TCEP and protease inhibitors. Hermes was eluted with a linear salt gradient (640 mM to 1 M NaCl). The fractions that contained the protein were concentrated and further purified at 4 °C on a preparative Superose 6 XK 16/70 size exclusion column (GE Healthcare) equilibrated with 25 mM HEPES.Na pH 7.5, 750 mM NaCl, 0.3 mM TCEP and protease inhibitors (see chromatogram in Fig. S1B). The fractions containing Hermes were then concentrated to be directly used or frozen in liquid nitrogen to be stored at −80 °C after the addition of glycerol to a final concentration of 15 %. Hermes eluted as a peak with a low molecular weight shoulder; we excluded the fractions corresponding to the shoulder (Fig. S1B).

### Preparation of the double-stranded DNA samples

All the oligonucleotides were purchased from Integrated DNA Technologies (IDT). The lyophilized DNAs were dissolved in 10 mM Tris-HCl pH 8.0 to a concentration of 1 mM or 1.5 mM for the unlabeled DNAs and to a concentration of 100 µM for the 6FAM-labelled DNAs. The double-stranded DNA samples were prepared by mixing stoichiometrically the complementary strands to a final concentration of 500 µM or 10 µM for the unlabeled and 6FAM-labeled DNAs. The mixes were heated to 95 °C for 10 minutes and slowly cooled to room temperature, and stored at −20 °C. All the DNAs used in this study are reported in Table S3.

### BED/DNA interaction assay monitored by size exclusion chromatography (SEC)

Purified BED was dialyzed overnight at 4 °C against 25 mM HEPES.Na pH 7.5, 150 mM NaCl, 0.3 mM TCEP and protease inhibitors (Roche) and subsequently concentrated (Vivaspin 20 3 kDa MWCO, GE Healthcare) to ~7 mg/mL. The samples (60 µL) were prepared by mixing the protein and the oligonucleotide from the 500 µM stock solution in various ratios (1:0, 0:1, 1:1, 2:1 and 3:1 DNA-to-protein ratio with 1 equivalent corresponding to 100 µM) and equilibrated on ice for at least 15 minutes. 10 µL of sample (10 µl injection loop loaded with 50 µL sample) were injected on a Superdex 75 PC 3.2/30 (GE Healthcare) analytical size exclusion column preequilibrated with the protein buffer. The UV absorbance at 280 and 260 nm were monitored to identify the different species.

### Hermes transposase oligomers stoichiometry as a function of its buffer composition determined by size exclusion chromatography (SEC)

Purified Hermes at 1 mg/mL in B750 buffer (750 mM NaCl 25 mM HEPES.Na pH 7.5, 0.3 mM TCEP and protease inhibitors) was either dialyzed overnight or 3 hours against buffer B500 (500 mM NaCl, 25 mM HEPES.Na pH 7.5 and 0.3 mM TCEP). The latter sample was then transferred in B250 (250 mM NaCl, 25 mM HEPES.Na pH 7.5 and 0.3 mM TCEP) either overnight or for 3 hours. The latter sample was finally transferred to B150 (150 mM NaCl, 25 mM HEPES.Na pH 7.5 and 0.3 mM TCEP) for 16 hours. The three samples were analyzed by SEC (Superose 6 30/100, GE Healthcare) in their respective buffer. The absorbance at 280 nm was monitored. To check whether the Hermes hexamer could be reversed to an octameric by buffer exchange, 1 mg/mL Hermes in B150 was dialyzed against B500 buffer overnight and injected on the SEC column preequilibrated with B500. The absorbance at 280 nm was monitored.

Calibration curves (MW as a function of the elution volume) in B500, B250 and B150 were obtained, respectively, using three proteins from the Sigma-Aldrich Gel Filtration Markers kit: thyroglobulin (669 kDa), apoferritin (443 kDa) and β-amylase (200 kDa).

### Hermes/DNA interaction assay monitored by size exclusion chromatography (SEC)

Purified Hermes was concentrated (Vivaspin 20 50 kDa MWCO, GE Healthcare) to ~3 mg/mL. The samples were prepared by mixing the protein and the oligonucleotide from the 500 µM stock solution in various ratios (1:0, 0:0.25, 1:0.25 and 1:0.5 protein-to-DNA ratio with 1 equivalent corresponding to 34 µM). The samples were successively dialyzed at 4 °C against 500, 250 mM and 150 mM NaCl containing buffers (25 mM HEPES.Na pH 7.5, 0.3 mM TCEP and protease inhibitors) for 2 hours, 4 hours and overnight, respectively. 100 µL of sample were injected on a Superose 6 30/100 (GE Healthcare) analytical SEC column preequilibrated with the last dialysis buffer. The UV absorbance at 280 and 260 nm were monitored to identify the different species.

### Mass photometry (MP) (Young et al. 2018)

#### Hermes transposase alone

The day before the experiment, three samples of frozen Hermes transposase (15 % glycerol, 25 mM HEPES.Na pH 7.5, 750 mM NaCl, 0.3 mM TCEP) were dialyzed for 2 hours against buffer B500 (500 mM NaCl, 25 mM HEPES.Na pH 7.5 and 0.3 mM TCEP). One sample was left in B500 to equilibrate overnight, while the other two were transferred in B250 (250 mM NaCl, 25 mM HEPES.Na pH 7.5 and 0.3 mM TCEP) for 3 hours. Finally, one sample was left overnight in B250, while the last sample was transferred to B150 (150 mM NaCl, 25 mM HEPES.Na pH 7.5 and 0.3 mM TCEP) for 16 hours. A volume increase as well as a light precipitation was observed as the NaCl concentration of the dialysis buffer decreased. The concentrations of the dialyzed samples were 10.0 µM (in B500), 6.5 µM (in B250) and 6.0 µM (in B150). Just before performing the MP experiments, the samples were diluted 10, 50 and 100 times in their respective buffers, and equilibrated for at least 10 minutes at room temperature. We used the detailed protocol published by the NHLBI biophysics facility with the silicon gasket applied on a glass coverslip as sample holder (Wu and Piszczek 2021). Data collection was performed on a G10 RefeynOne mass photometer. 10 µL of buffer was used to optimize the focus, then 10 µL of sample was mixed to the buffer drop for data collection (1 min acquisition). The concentrations that gave the best signal-to-noise were 50 nM, 65 nM and 30 nM in B500, B250 and B150, respectively. The movie frames were processed with the built-in software DiscoverMP. The contrast values were converted to masses using a calibration curve obtained with BSA (monomer and dimer), alcohol dehydrogenase (monomer and dimer), ovalbumin, and thyroglobulin in PBS buffer. The mass distributions were fitted using a Gaussian distribution model implemented into DiscoverMP. The theoretical mass of a Hermes monomer, dimer, hexamer and octamer are 70.1, 140.2, 420.7, and 561.0 kDa, respectively.

#### Transposase/DNA complexes

A typical sample (100 µL) was composed of 10 µM Hermes mixed with 2.5, 5 or 10 µM DNA in 25 mM HEPES.Na pH 7.5, 750 mM NaCl and 0.3 mM TCEP. The samples were successively dialyzed as described for the transposase alone to reach the equilibrium in B150. Just before the experiments, the samples were diluted 50 and 100 times and equilibrated for at least 10 minutes at room temperature. The same experimental procedure was used as described for the apo protein. The composition of the protein/DNA complexes (P1 and P2 masses) were determined by subtracting the mass of either hexameric Hermes or octameric Hermes, and the remaining mass was then divided by the mass of the given DNA. The theoretical MW of the Hermes hexamer and octamer are 420.7 and 561.0 kDa, respectively. The theoretical MWs of the LE-TIR, LE-TIR+13, LE-TIR+30 and 8+LE-TIR+30 DNAs are 10.1, 18.1, 32.2 and 33.2 kDa, respectively.

### Crystallization, X-ray diffraction data collection, structure determination and model refinement of the BED/LE11-27 complex

After purification, BED was dialyzed overnight at 4 °C against 25 mM HEPES.Na pH 7.5, 100 mM NaCl, 0.3 mM TCEP and subsequently concentrated (Vivaspin 20 3 kDa MWCO, GE Healthcare) to ~10 mg/mL. The LE11-27 DNA was dialyzed against the same buffer and concentrated to ~ 2.5 mM (Vivaspin 500, 3 kDa MWCO, Merck). The complex was formed by mixing the protein and the DNA in a 2:1 protein-to-DNA ratio (final concentration of 800 µM and 400 µM, respectively). Crystals were grown at 20 °C by the hanging drop method. 1.7 µL of sample were mixed with 2.3 µL of crystallization solution composed of 100 mM Bis-Tris pH 6.5 (Hampton Research) and 25 % PEG 4000 (Hampton Research). Crystals grew over 16 days to a size of ~ 0.3 x 0.3 x 0.3 mm and were cryoprotected by a quick transfer to a stabilizing solution at 20 % ethylene glycol, 12.5 mM HEPES.Na pH 7.5, 50 mM NaCl, 25 mM Bis-Tris pH 6.5, 12.5 % PEG 4000 prior freezing in liquid nitrogen. The X-ray diffraction data were collected at the Advanced Photon Source beamline 22-ID, operated by SER-CAT on an Eiger X16M detector. Three anomalous diffraction data sets were collected on the same crystal around the zinc absorption edge, peak and remote wavelengths (9661 eV and 1.28335 Å; 9665 eV and 1.28282 Å; 10000 eV and 1.23984 Å, respectively). The diffraction data were integrated and scaled with XDS and XSCALE (Kabsch 2010). Initial experimental electron density maps were computed with Autosol from the Phenix package. (Adams et al. 2010). The maps were improved by incorporating the anomalous signal of the DNA backbone’s 8 P atoms and the 4 S atoms from the protein using an additional data set collected at 8.03 keV using a rotating anode source equipped with an Eiger 4M detector. Phase calculations were performed by using Sharp (De La Fortelle and Bricogne 1997). The density-modified map enabled to build an initial model in COOT (Emsley et al. 2010). The “edge” diffraction data processed as a non-anomalous dataset (Friedel law considered as true) and treated for anisotropy (http://staraniso.globalphasing.org/cgi-bin/staraniso.cgi) was used to further improve and refine the model in Phenix (Adams et al. 2010) and Buster (Global Phasing Limited) (Bricogne and IUCr 1993) at 2.5 Å resolution. Detailed crystallographic statistics are in Table S1.

### Cryo-EM specimen preparation, cryo-EM data collection, single-particle analysis, and model building

Frozen or freshly purified Hermes transposase was mixed in an 8:2 protein-to-DNA ratio with either 8+LE-TIR+30 or 8+RE-TIR+30 in a 250 µL sample (1 equivalent was equal to 5 µM). The samples were dialyzed against a series of buffers as described for the SEC interaction assay with the final buffer composed of 25 mM HEPES.Na pH 7.5, 150 mM NaCl, 0.3 mM TCEP and protease inhibitors (Roche). The samples were concentrated to 100 µL and purified by SEC (Superose 6 30/100, GE Healthcare) at 4 °C. The fraction with the highest absorbance was concentrated to reach the absorbance of ~0.60 at 260 nm and ~0.5 at 280 nm. The protein concentration was estimated to be ~0.2 mg/mL by comparing the SDS-PAGE band intensity of the cryo-EM sample with that of a dilution series of apo-Hermes.

The samples were frozen on copper or gold grids with holey carbon film coated with a 2 nm continuous carbon film (Quantifoil R1.2/1.3 ultrathin carbon, 300 mesh) freshly glow discharged for 20 s at 15 mA (PELCO easiGlow). A Vitrobot Mark IV (FEI) rapid plunging device was used for the specimen preparation. The Vitrobot chamber was at room temperature and the relative humidity set at 100 %. Three µL of samples were applied on the grid and after 10 s wait, the excess sample was blotted for 3 or 4 s (force 1) and flash frozen in liquid ethane cooled by liquid nitrogen. We used carbon coated grids to partially prevent the complex from falling apart due to denaturation at the air/water interface during the specimen preparation (D’Imprima et al. 2019).

For the LE-LE transpososome, we collected ~9600 movies into two sessions on a 200 kV Glacios TEM (FEI) equipped with a K3 direct electron detector camera (Gatan). The movies were recorded in super resolution counting mode at a nominal magnification of 130,000, corresponding to a calibrated super resolution pixel size of 0.58 Å per pixel with a defocus range from −1 to −2.5 µm. The acquisition was supervised by the semi-automated program SerialEM (Mastronarde 2005). The dose rate on the camera was set at 15 electron per physical pixel per second. The total exposure time for each movie was 2 s with a total exposure dose of 22.3 e^−^/Å^2^ (1.39 e^−^/Å^2^ per frame). Each movie was composed of 16 frames, with 125 ms per frame.

For the RE-RE transpososome ~9500 movies were recorded on a 300 kV Titan Krios TEM (FEI) equipped with a K3 camera. The movies were recorded in super resolution counting mode at a nominal magnification of 130,000 corresponding to a calibrated super resolution pixel size of 0.43 Å per pixel with a defocus range from −1 to −2.5 µm. The acquisition was supervised by the semi-automated program SerialEM (Mastronarde 2005). The dose rate on the camera was set at 22 electron per physical pixel per second. The total exposure time for each movie was 1.66 s with a total exposure dose of 48.7 e^−^/Å^2^ (2.21 e^−^/Å^2^ per frame). Each movie was composed of 22 frames, with 75 ms per frame.

The single particle analyses were performed with RELION 3.1 (Scheres 2012a, 2012b; Zivanov et al. 2018) ran on the NIH HPC Biowulf cluster (http://hpc.nih.gov). The processing of each dataset followed the same protocol until stated differently. The processing workflows are presented in Fig. S6, S7 and S8. UCSF Chimera (Pettersen et al. 2004) was used to visualize the maps. The figures of cryo-EM map and models were prepared with Chimera or ChimeraX (Goddard et al. 2018) and PyMol (http://www.pymol.org).

The movies were motion corrected in RELION3.1 and binned by a factor 2, resulting in a pixel size of 1.16 Å (Glacios data) and 0.86 Å (Krios data), respectively, for further processing. The contrast transfer function (CTF) parameters were estimated with gctf1.06 (Zhang 2016). The program crYOLO (Wagner et al. 2019) was used for particle picking with a general model on motion corrected movies (~3.85 million and ~2.92 million initial particles for LE-LE and RE-RE transpososome, respectively). The best particles were selected by several rounds of 2D classifications. The particle stacks of the LE-LE transpososome were joined and ran through a last 2D classification. The resulting particle stack (~641,300 particles and ~186,493 particles for LE-LE and RE-RE transpososome, respectively) was used to generate a 3D initial model that was used as reference for a 3D classification job (six classes). The best classes were used for gold-standard refinement (class #4 and #6 of the LE-LE transpososome and class #2 of the RE-RE transpososome in Fig. S6 and Fig. S8, respectively). For both samples, the equatorial Hermes dimers (B and D) were poorly resolved, and we opted for a partial signal subtraction strategy to exclude them from the experimental images and improve the alignment of the core of the complex.

For the LE-LE transpososome a “consensus” gold-standard refinement was performed with the joined particles from 3D classes #4, #5 and #6 (~359,800 particles), followed by per-particle CTF refinement and Bayesian polishing. A mask excluding the equatorial Hermes dimers was generated for partial signal subtraction on the same stack of particles. The resulting “edited” particles were subjected to two rounds of focused 3D classifications without orientational search. Gold-standard refinement with imposed C2-symmetry was performed with the particles from the best classes (classes 2, 3 and 4, ~143,400 particles), i.e., classes that showed better density for the BED domains. Another round of CTF refinement and Bayesian polishing was performed using the corresponding “whole” particles. Finally, the shiny particles were signal subtracted and gold-standard refined. The resolution of the final map was 4.64 Å at 0.143 Fourier Shell Correlation (FSC) threshold.

For the RE-RE transpososome, the 3D class #2 was gold-standard refined to generate a mask that only covered the core of the transpososome, leaving out the equatorial Hermes dimers. Partial signal subtraction was performed on the particle stack of class #2 (53,656 particles). The resulting “edited” particles were subjected to focused refinement with local orientational search. The resolution of the final map was 5.31 Å (0.143 FSC threshold).

Using UCSF Chimera (Pettersen et al. 2004), we built in an approximative atomic model into the LE-LE transpososome map by rigid-body fitting two crystal structures of the N-truncated Hermes dimer bound to two nicked TIRs (PDB:6DX0) (Hickman et al. 2018) as well as four BED/LE11-27 crystal structures (PDB: 8EB5). The 3’-end of the LE-DNAs were modelled using the DNA from the crystal structure of the truncated Hermes dimer bound to two cleaved TIRs (PDB: 4D1Q) (Hickman et al. 2014). A protomer of the crystal structure of the apo N-truncated Hermes (PDB: 2BW3) poorly fit the density of the protomers A2 and C2; the holo structure was the best fit. The DNA base pairs and the BED domains that laid out of the density were removed. The contiguous pieces of DNAs were then merged as one dsDNA, and sequence corrected in COOT (Emsley et al. 2010) to generate two LE-DNAs. The BED domains of protomers A1, A2, C1 and C2 were linked to the N-truncated Hermes structures in COOT as well. Using COOT 0.9.6.2-pre EL (Casañal, Lohkamp, and Emsley 2020), each chain was globally real space refined. In addition, self-local distance restraints as well as B-form restraints were applied on the DNA chains. The resulting model was real-space refined against the cryo-EM map in Phenix 1.19.2 (cryo-EM module) to finalize the flexible fitting and correct some geometries. The refinement statistics are summarized in Table S2.

### Electrophoretic mobility shift assay (EMSA)

#### BED/DNA EMSAs

The samples were prepared by mixing 0.9 µM of cold DNA in B150 (25 mM HEPES.Na pH 7.5, 150 mM NaCl, and 0.3 mM TCEP) with 0.1 µM of 6FAM-labelled DNA in B150 (5’-6FAM top strand), 0.5 µM of ran17 DNA, various amount (0 to 10 µM) of BED protein in B150 and loading dye (0.5X TBE, 10 % glycerol and bromophenol blue). The samples were equilibrated for 1 hour at 4 °C, while the PAGE gels were pre-run at 80 V (15 % polyacrylamide, 0.5X TBE, 1 mm gels). The samples were equilibrated for 15 minutes at room temperature prior being loaded (10 µL) onto the gel. The PAGE was run for 4 hours at 80 V in an ice box. The gels were scanned with a Typhoon FLA7000 (GE Healthcare) fluorescence imager. The gels whose fluorescence intensity was meant to be extracted were underexposed to avoid the saturation of the detector. We used ImageJ to extract the intensity of the bands (Schneider, Rasband, and Eliceiri 2012). The band intensities were transformed as percentage of DNA bound and plotted against the concentration of BED. The fitting of the data fitting was performed in GraphPad Prism 9.3.1.

### In cell transposition assay

The helper plasmid pFV4a-Hermes and the donor plasmids pHermesWT-CMVpuro, pHermes2LE-CMVpuro, pHermes2RE-CMVpuro, pHermes2LEmut-CMVpuro, pHermesLE50Scr-CMVpuro and pHermesLE140Scr-CMVpuro were ordered from GenScript using the plasmids pFHelR and pHelR-CMV-puro from Grabundzija et al. (2018) as backbones (Grabundzija, Hickman, and Dyda 2018). The rest of the donor plasmids were obtained by deletion mutagenesis of pHermesWT-CMVpuro (Makarova, Kamberov, and Margolis 2000). The sequences of the transposon ends are reported in Table S4. The helper plasmid contains the gene of the Hermes transposase, and the donor plasmid carries the puromycin resistance gene flanked by the *Hermes* LE and RE and variants to form the puro^R^ transposon.

HEK293T cells (0.5 x 10^6^) were seeded onto six-well plates 1 day before transfection. The cells were transfected with 1 µg of helper plasmid and 0.5 µg of the donor plasmid. All the transfections were performed with Lipofectamine 3000 (Thermo Fisher) according to manufacturer’s protocol. Two days post-transfection, the cells was replated onto 100 mm dishes at a 50-fold dilution (i.e., 2 % of the cells) and selected for transposon integration with 2 µg/mL puromycin. The selection medium was changed every 3 days. After 8-12 days, the cell colonies were fixed directly into their dish for 20 minutes with 4 % formaldehyde in PBS and subsequently stained overnight with 1 % methylene blue in PBS for counting. All experiments were independently replicated at least 4 times. Negative control experiments were performed by replacing the helper plasmid with pUC19. The *piggyBac* system was used as a procedure control.

All statistical analyses were performed using GraphPad Prism 9.3.1. Two-tailed Student’s unpaired t-tests were used to compare means between experimental samples and their corresponding pUC19 controls and means between some experimental samples and pHermesWT-CMVpuro. The bars correspond to the standard deviation centered on the mean.

## Data Availability

The crystal structure of the Hermes transposase BED domain bound to the Hermes transposon LE quasi-palindrome LE-STR1-STR2 was deposited in the Protein Data Bank (PDB) under the accession code 8EB5.

The cryo-EM map of the core of the LE-LE *Hermes* transpososome was deposited in the Electron Microscopy Data Bank (EMBD) under the accession code 28034 and its related structure model was deposited in the PDB with the code 8EDG.

## Acknowledgements

This work was supported by the Intramural Program of the National Institute of Diabetes and Digestive and Kidney Diseases, National Institutes of Health (NIDDK/NIH) under Grant DK036153-16. X-ray diffraction data were collected at the Advanced Photon Source, Argonne National Laboratory on the 22-ID beamline operated by Southeast Regional Collaborative Access Team (SER-CAT). SER-CAT is supported by its member institutions (www.ser-cat.org/members.html), and equipment grants (S10_RR25528 and S10_RR028976) from the National Institutes of Health. The use of the Advanced Photon Source was supported by the U. S. Department of Energy, Office of Science, Office of Basic Energy Sciences, under Contract No. W-31-109-Eng-38. The computations for the cryo-EM single particle analysis were carried out using the High-Performance Computing Systems at the NIH.

The authors thank Greg Piszczek and Di Wu of the Biophysics Core Facility at the National Heart, Lung, and Blood Institute (NHLBI/NIH) for the access to the mass photometer. The authors would like to also thank Huaibin Wang and Yanxiang Cui for their help with cryo-EM data collection at the Multi Institute Cryo-EM Facility (MICEF), NIDDK/NIH. The authors are also thankful to Istvan Botos (NIDDK/NIH) for on-site computational support.

## Author contributions

LL performed most of the experiments in the study except for the in-cell transposition assays on the truncated LE that were carried out by CF. LL analyzed all the data generated in the study. LL and FD analyzed the X-ray data. LL prepared all the samples used in this study including the crystals and the cryo-EM specimens. ABH initiated the project. FD supervised the study. All authors participated in manuscript preparation.

## Supplementary information

**Figure S1:**
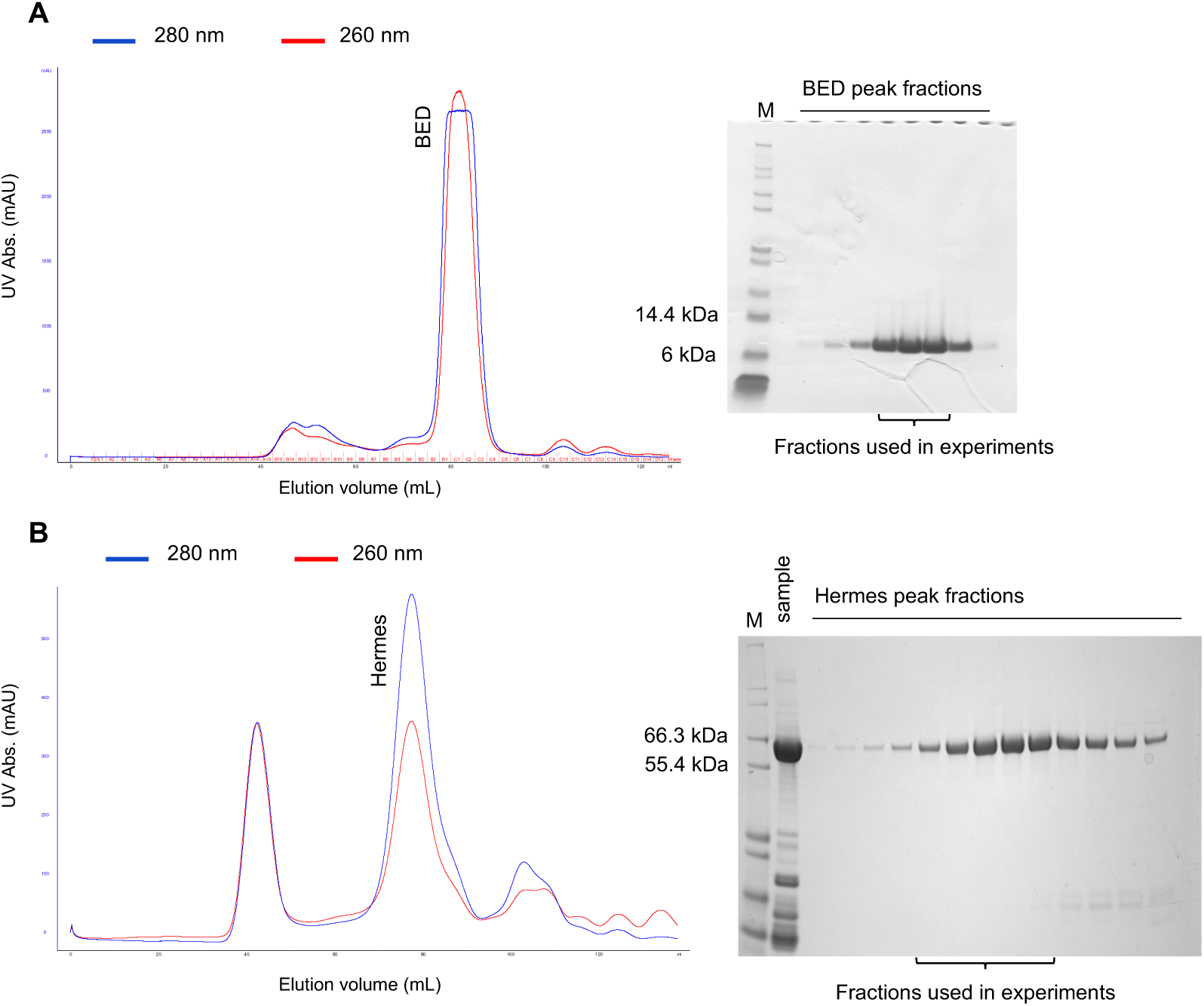
Final purification step (size exclusion chromatography or SEC) of the Hermes BED domain (A) and of the full-length Hermes transposase (B). The SEC chromatograms are on the left and the SDS-PAGE analysis of the collected fractions are on the right. M: protein size marker (Mark12, Novex).

**Figure S2:**
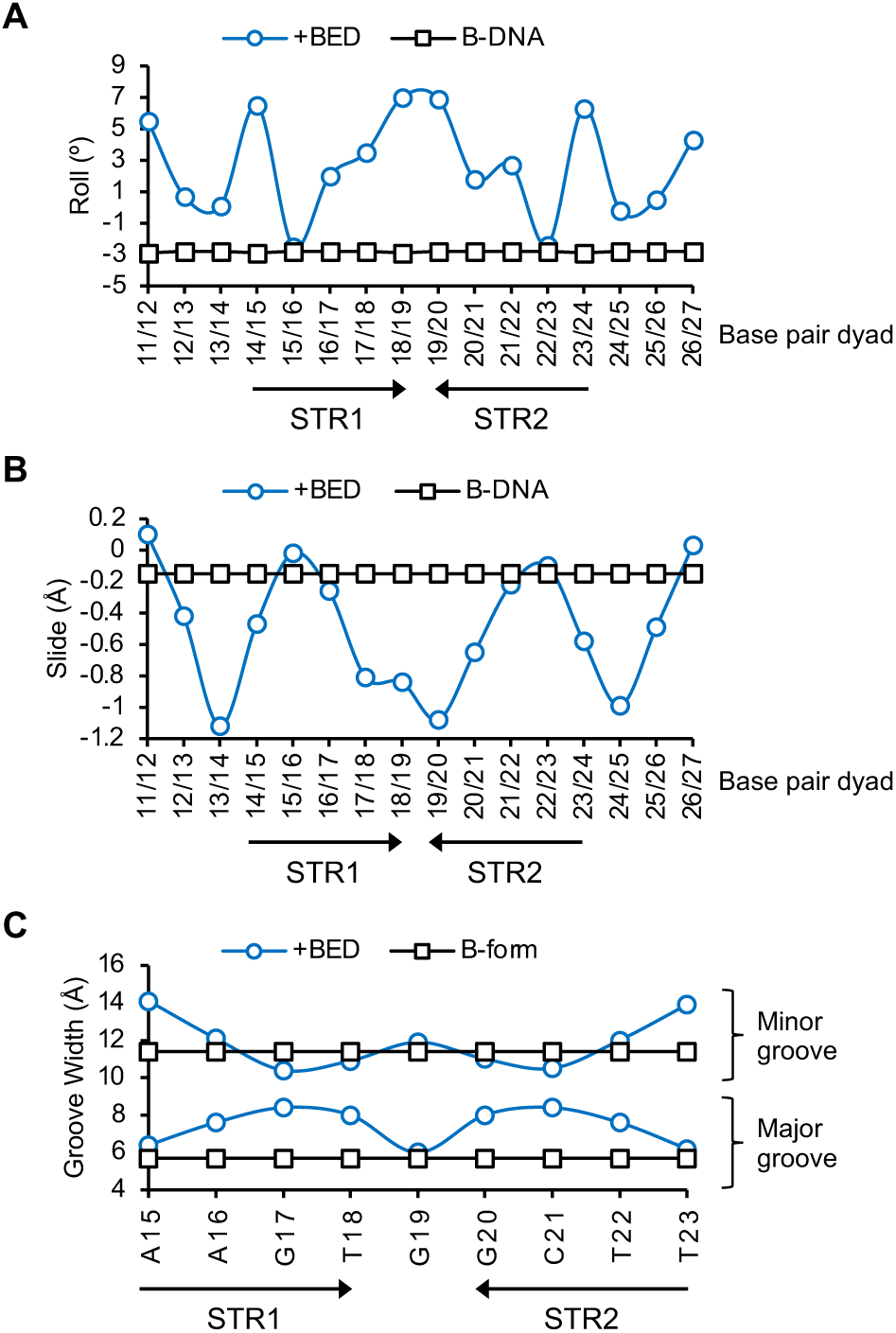
The roll (A), slide (B) and groove parameters (C) of the LE11-27 DNA bound to Hermes BED domains (crystal structure from Fig. 2 (PDB: 8EB5)) compared to that of measured in its B-form.

**Figure S3:**
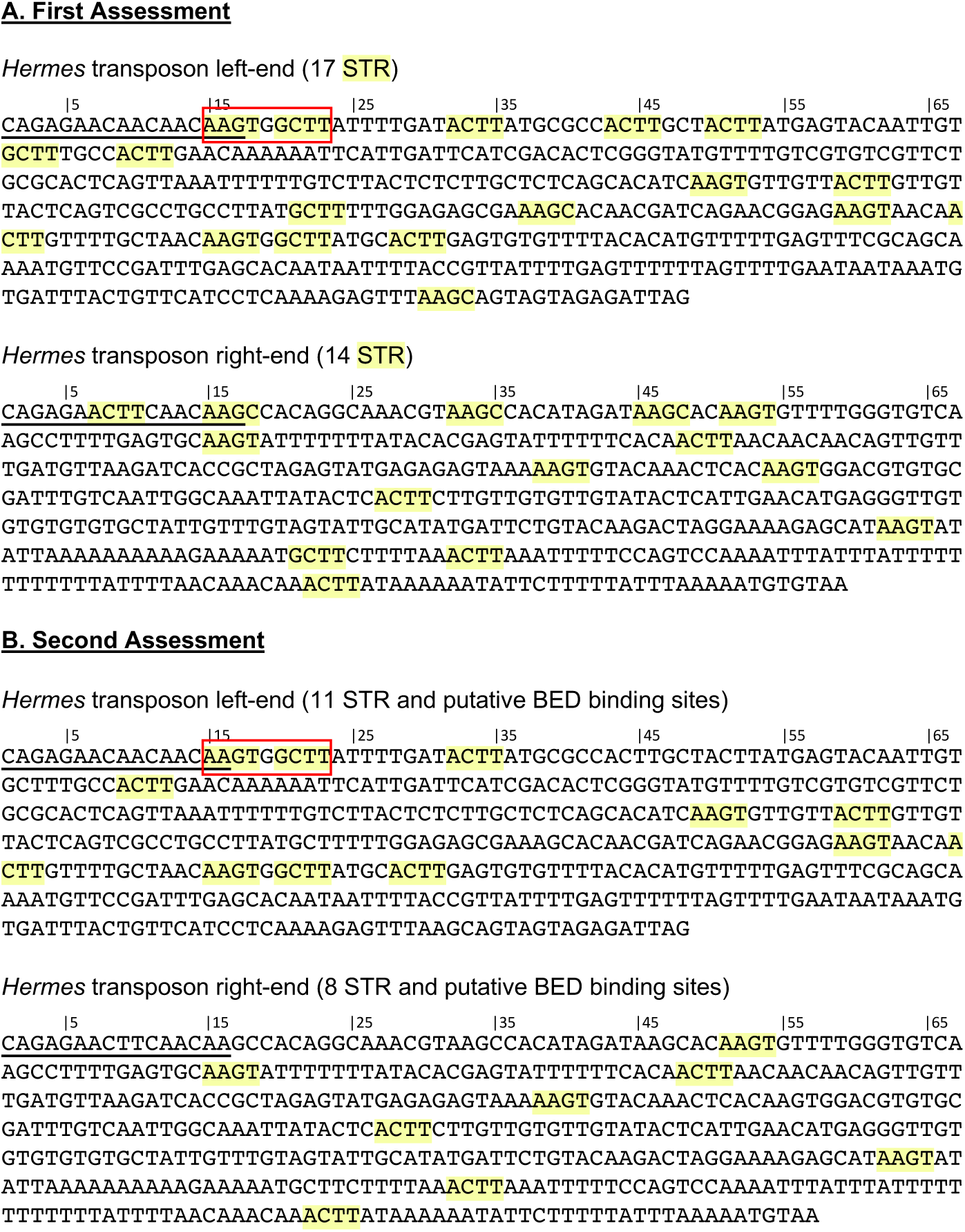
Mapping on the *Hermes* transposon ends of the subterminal repeats (STR) proposed as putative BED binding sites. A) Mapping based on the definition of the STR as 5’-AAGY-3’ with Y being T or C. B) Mapping based on a finer definition of the BED binding site that corresponds to 5’-AAGT-3’ followed by an AT-rich sequence or in the special situation of the quasi-palindrome 5’-AAGTGGCTT-3’. The LE-STR1-STR2 quasi palindrome that cooperatively interact with two BED domains is highlighted by a red box.

### The stoichiometry of the Hermes transposase

During the purification of the Hermes transposase, we observed heterogeneity and unusual behavior when the samples were analyzed by size exclusion chromatography (SEC) at different ionic strengths. As shown in Fig. S4A (same as Fig. 3H) the purified transposase itself eluted as two populations, P1 and P2, where the relative ratio between the two elution peaks varied as a function of the salt concentration in the buffer, with the higher concentration favoring P1 and lower concentrations promoting P2. At the protein concentration used, lowering the ionic strength caused precipitation and the data suggested that P2 stays more soluble at lower ionic strengths relative to P1. When the ionic strength was dropped, the relative proportion of P1 to P2 did not change and stayed the same when it was raised again (Fig. S4A), ruling out a simple and reversible interconversion process.

According to the standard curve obtained at the same elution conditions, P1 and P2 had an apparent molecular weight (MW) of 558 kDa and 434 kDa, respectively, corresponding closely to an octamer (theoretical MW of 561 kDa) and a hexamer (theoretical MW 421 kDa), respectively. To confirm that the two populations observed by SEC were indeed the result of a mass difference and not conformational variants of the same complex, we performed mass photometry (MP) (Sonn-Segev et al. 2020; Young et al. 2018). The MP histograms obtained in buffers with 500, 250 and 150 mM NaCl are shown in Fig. S4C. Four distinct masses were detected in all experiments and were attributed to the Hermes monomer (P4 mass average: 83 kDa for a theoretical MW of 71 kDa), dimer (P3 mass average: 133 kDa for a theoretical MW of 140 kDa), hexamer (P2 mass average: 425 kDa) and octamer (P1 mass average: 571 kDa), respectively. Although we never observed Hermes monomers and dimers by SEC, they were present in solution according to the MP data, and together roughly represented 20-30% of the total Hermes particles detected. We note that at high ionic strength more octamers were counted, whereas at lower ionic strength more hexamers were counted. These observations confirmed the initial results of the SEC experiment. Through a combination of SEC and mass photometry (MP) analysis, we established that the Hermes transposase exists in solution as two oligomeric states composed of six or eight monomers.

---

**Figure S4:**
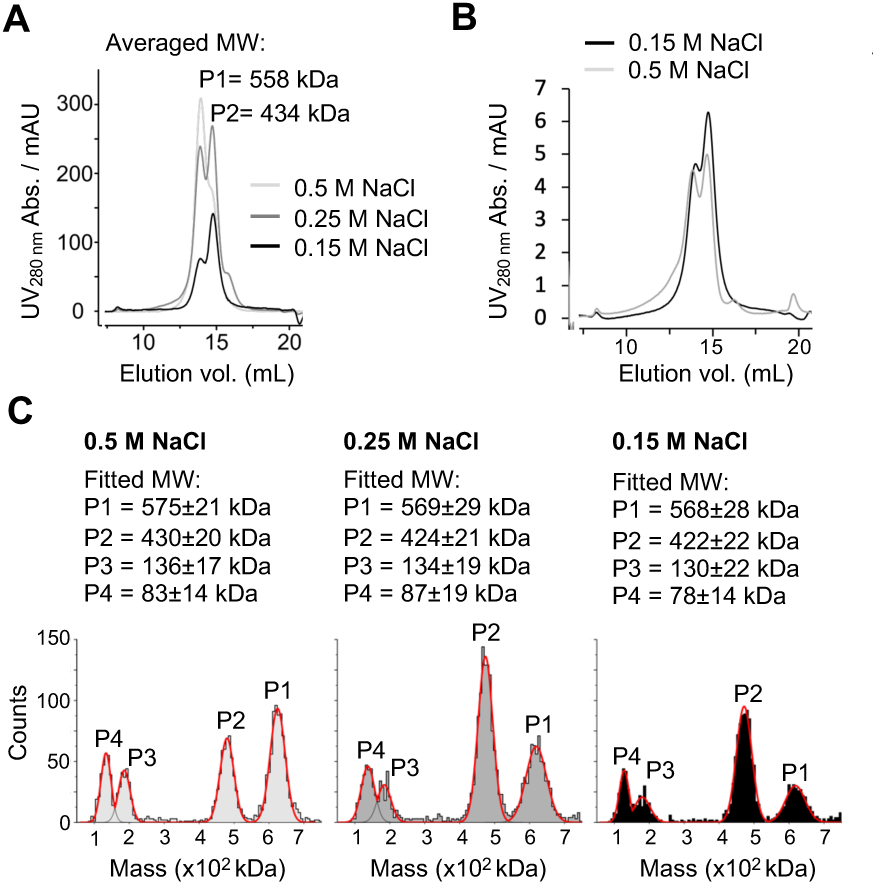
Stoichiometry of the Hermes transposase assembly. A) Size exclusion chromatogram of purified Hermes (in 0.75 M NaCl) dialyzed against three different elution buffers containing 0.5 M (light grey), 0.25 M (grey) and 0.15 M (black) NaCl. The apparent molecular weight (MW) derived from a standard curve for P1 and P2 is reported. B) Size exclusion chromatogram of purified Hermes (in 0.75 M NaCl) buffer exchanged to lower the salt concentration to 150 mM NaCl (black) then re-buffered in a 500 mM NaCl buffer (grey). C) Mass photometry histograms of similar samples as in A). The mass of P1, P2, P3 and P4 were obtained from a standard curve, and the reported errors correspond to the fitting error to a Gaussian distribution model. The theoretical mass of a Hermes monomer, dimer, hexamer and octamer are 70.1, 140.2, 420.7, and 561.0 kDa, respectively.

**Figure S5:**
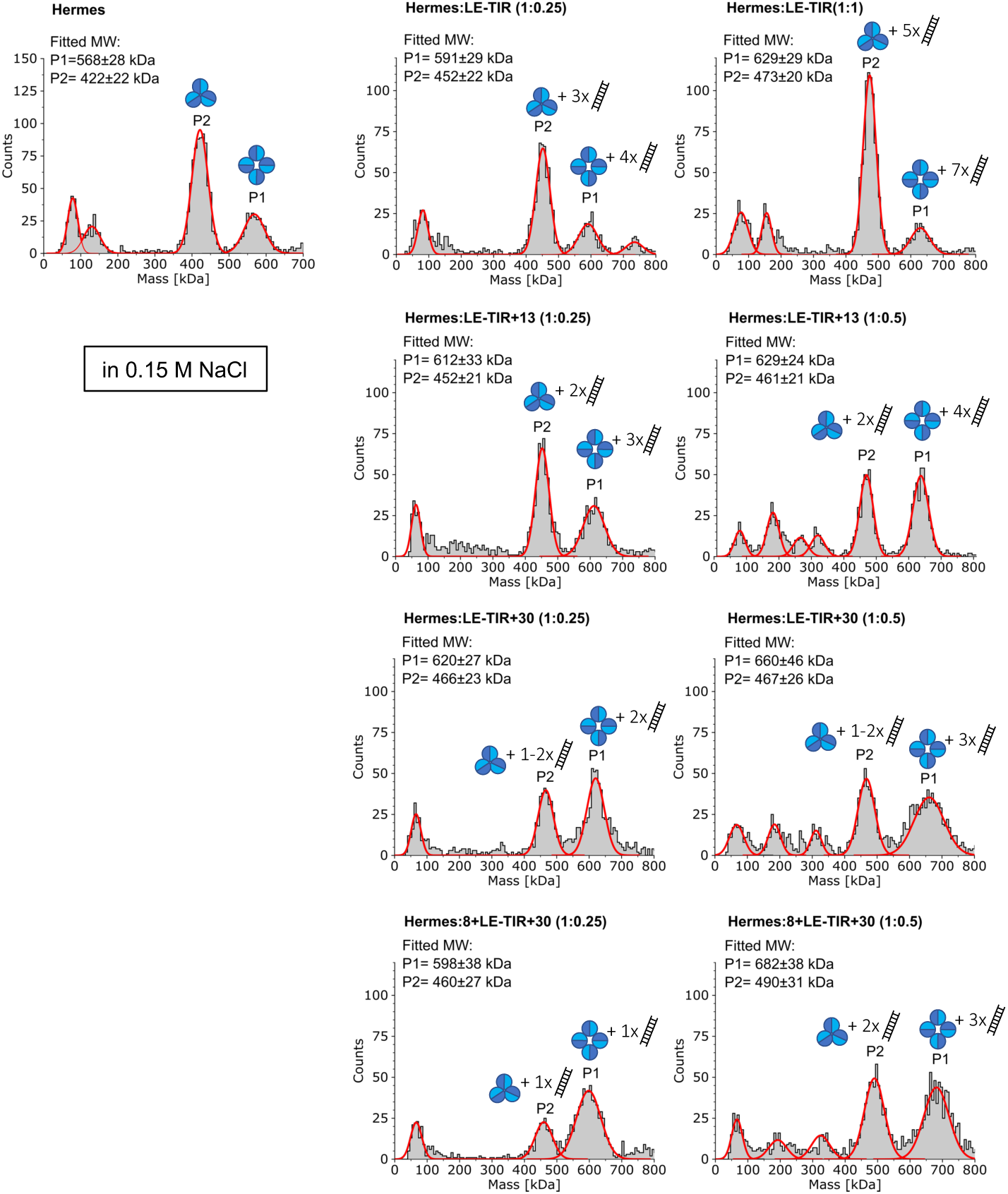
Characterization of the molecular composition of the Hermes transposase in complex with different Left End (LE) DNAs by mass photometry. The fitting masses of P1 and P2 were reported as well as their molecular composition. The hexameric and octameric forms of Hermes are represented by 3 and 4 blue circles, respectively. The DNAs are schemed as ladders. The theoretical MW of a Hermes hexamer and octamer are 421 and 561 kDa, respectively. The theoretical MWs of the LE-TIR, LE-TIR+13, LE-TIR+30 and 8+LE-TIR+30 are 10, 18, 32 and 33 kDa, respectively.

**Figure S6.**
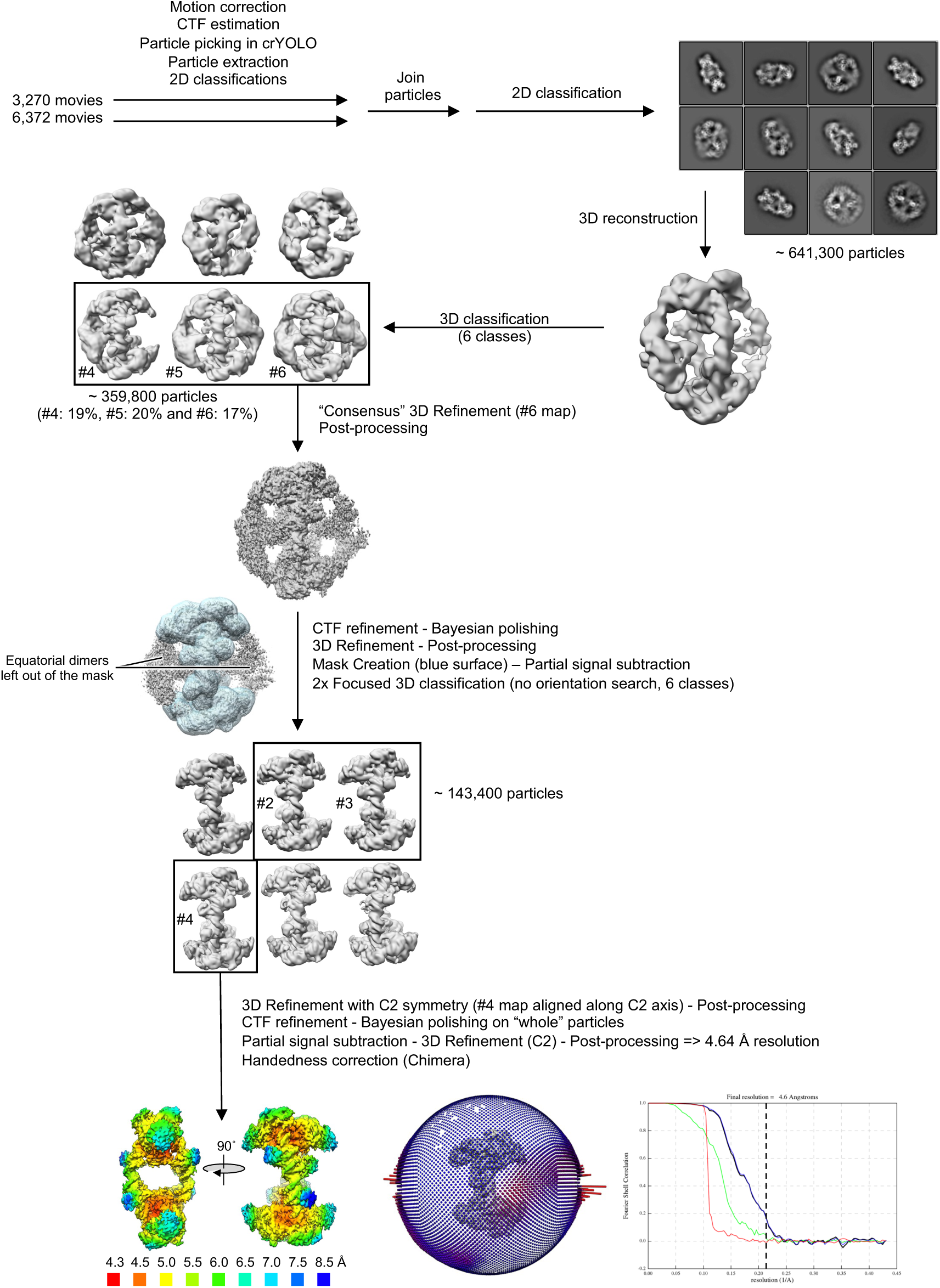
Cryo-electron microscopy single particle analysis workflow of the LE-LE transpososome in RELION.

**Figure S7:**
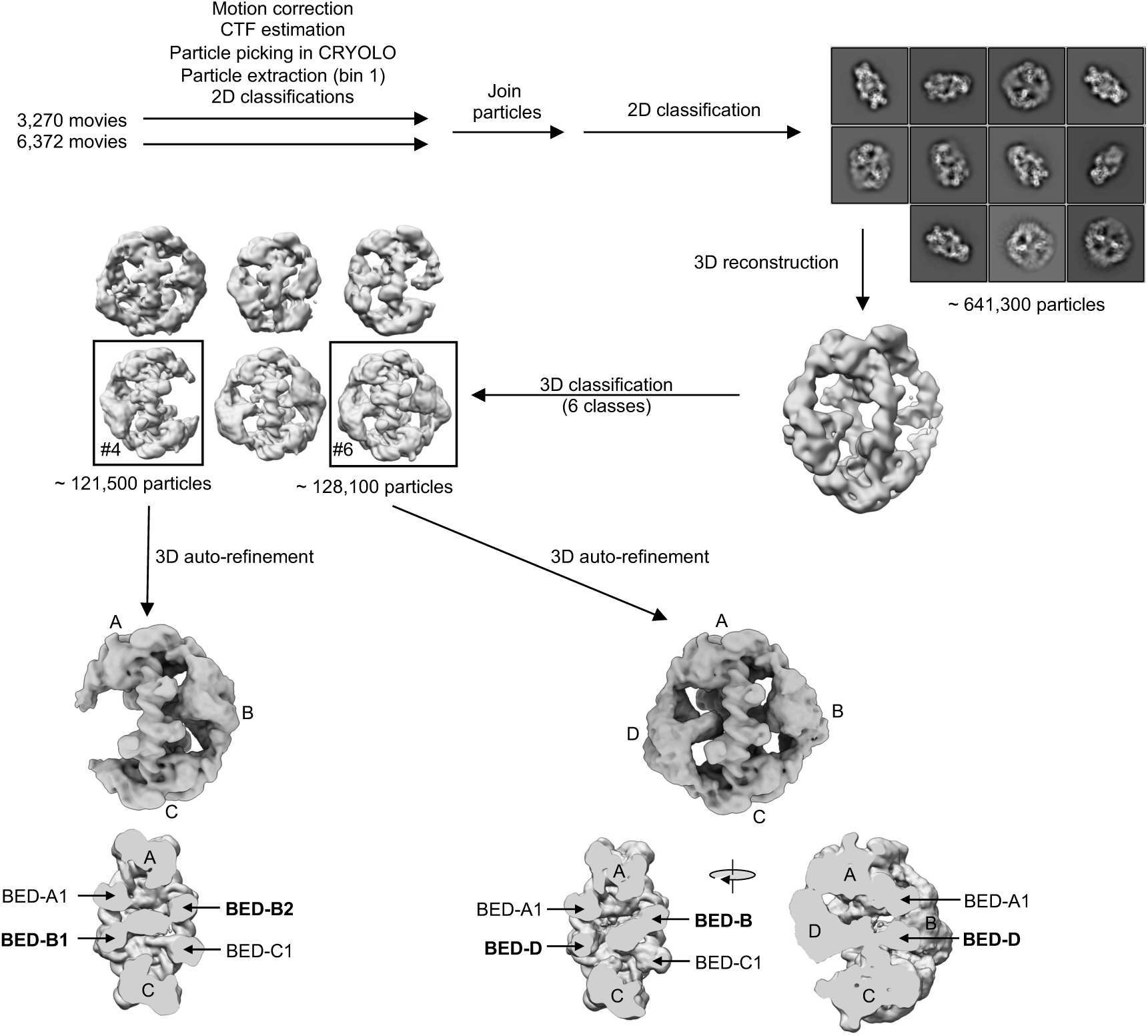
Identification of the origin of the BED domains interacting with the LE-STR2. In this branching of the single particle analysis of the cryo-EM data of the LE/LE *Hermes* transpososome, the equatorial Hermes units B and D were not masked-out.

**Figure S8.**
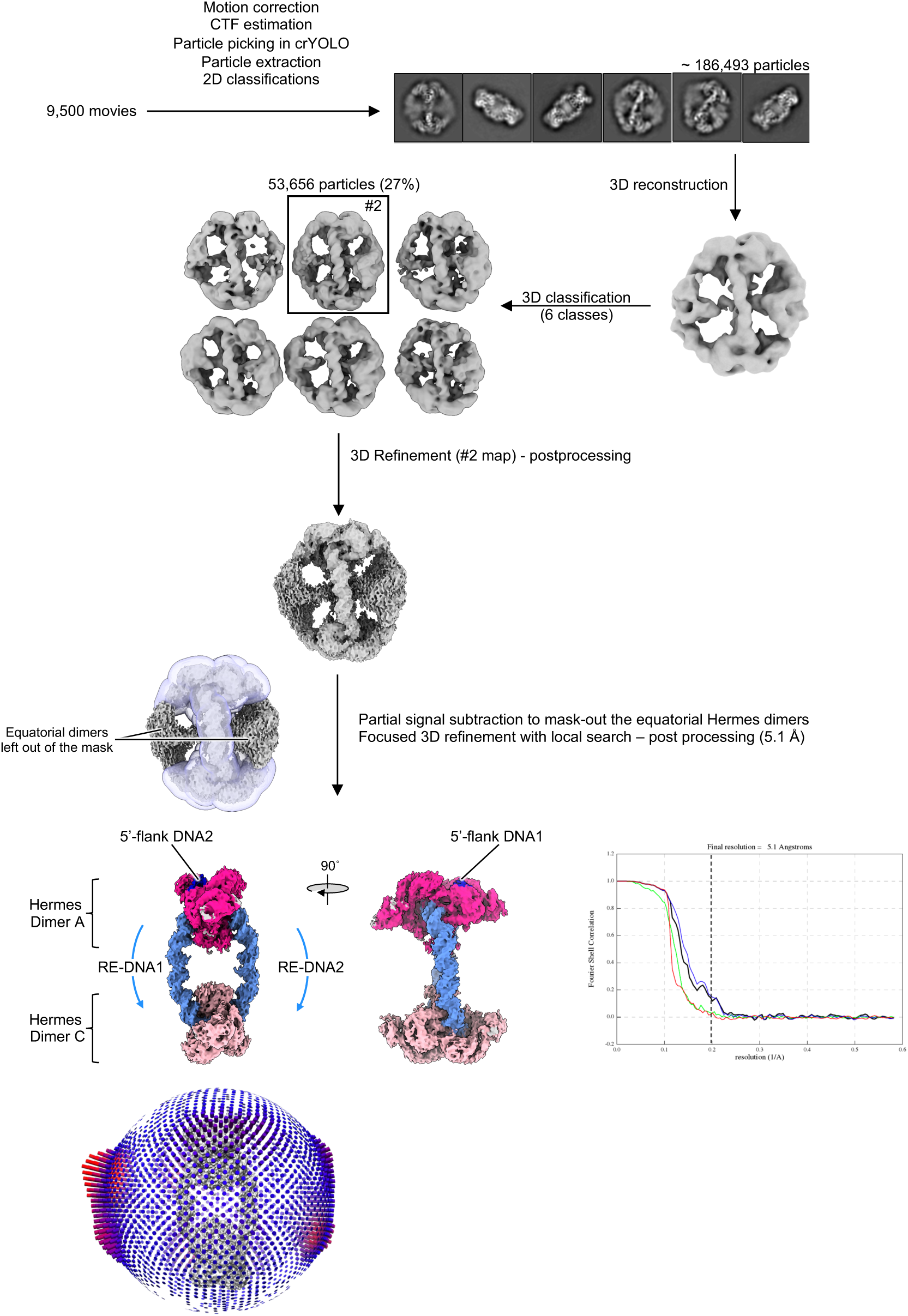
Cryo-electron microscopy single particle analysis workflow of the RE/RE *Hermes* transpososome in RELION.

**Figure S9:**
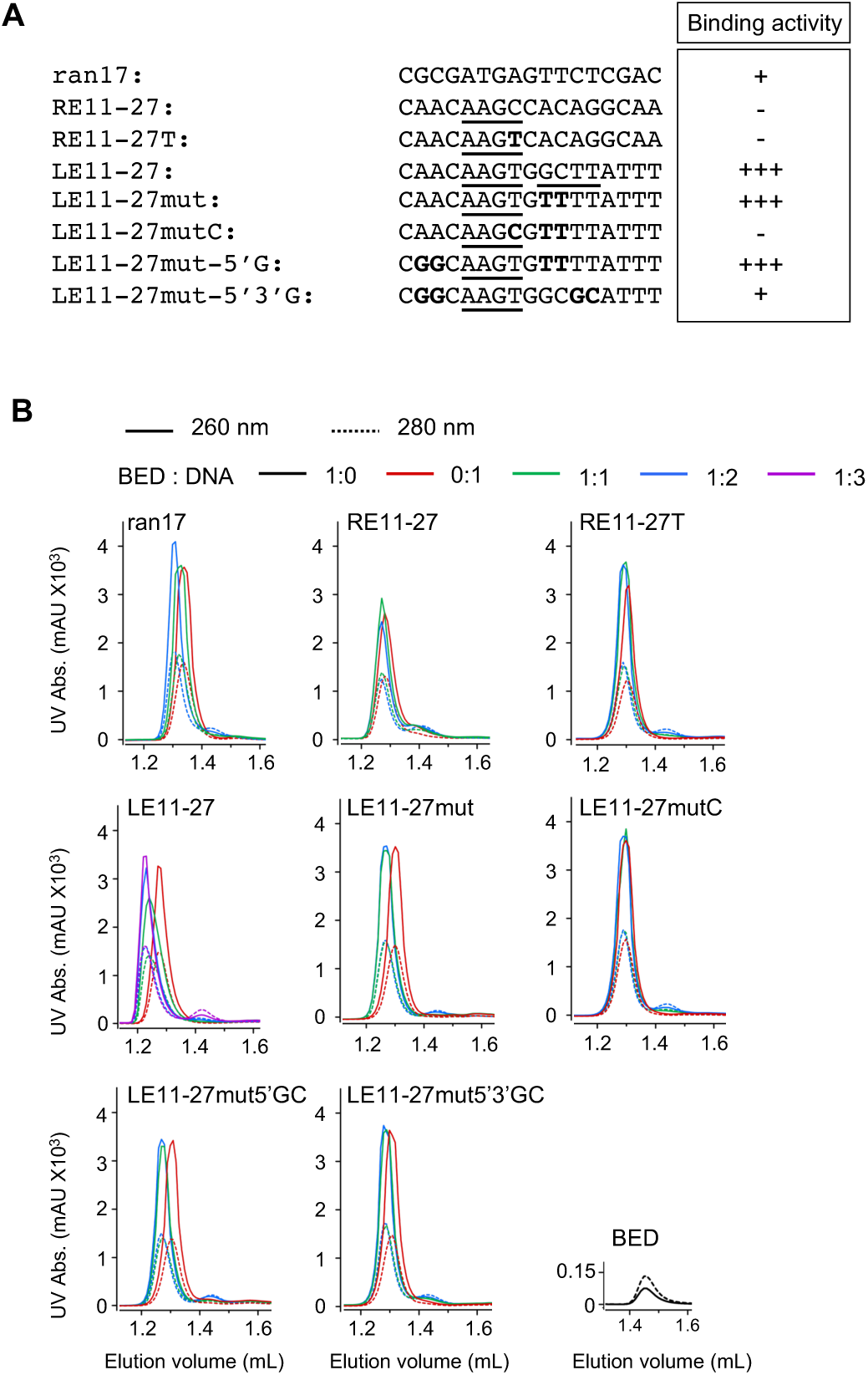
The minimal BED binding site. A) Top strand of the Left End (LE) and Right End (RE) DNAs used in the interaction assay. The 5’-AAGY-3’ (with Y = C or T) motifs are underlined, and the mutations are in bold. The table summarizes the binding activity of the Hermes BED domain for each oligo. B) Size exclusion chromatograms of the BED/DNA mixes. The BED-to-DNA ratios of the samples are reported on top.

**Figure S10.**
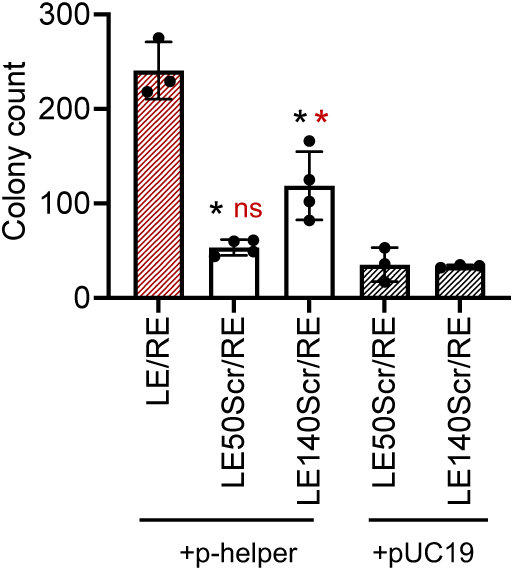
Hermes transposition assay in HEK293T cells with partially scrambled Left End (LEScr) (related to Fig. 7). In the plasmid donors LE50Scr/RE and LE140Scr/RE the *Hermes* LE was scrambled after the base pairs 50 and 140, respectively (instead of being truncated as in the assay reported in Fig. 7). Thus, the distance between the LE-TIR and the CMV promoter remains constant. Each circle represents one independent data point, with n = 3 or 4. The mean value is plotted as a column and the standard deviation as a bar. The statistical t-test was applied to determine the significance of the results compared to the LE/RE experiment (black star) or the appropriate negative control (+pUC19) (red star or red ns). *: p ≤ 0.01, ns: not significant.

**Table S1.** Crystallographic statistics

|  | Zn-MAD |  |  | Rotating Anode |
| --- | --- | --- | --- | --- |
|  | "Edge" | "Peak" | "Remote" |  |
| Wavelength (Å) | 1.28335 | 1.28282 | 1.23984 | 1.54184 |
| Resolution range (Å) | 27.6-2.5 (2.56-2.5) | 27.6-3.0 | 27.6-3.0 | 27.6-2.7 (2.77-2.7) |
| Space group | P6<sub>5</sub>22 | P6<sub>5</sub>22 | P6<sub>5</sub>22 | P6<sub>5</sub>22 |
| Unit cell dimensions<br>a, b, c (Å) and $\alpha$, $\beta$, $\gamma$ (°) | 55.3, 55.3, 212.6<br>90, 90, 120 | 55.3, 55.3, 212.6<br>90, 90, 120 | 55.3, 55.3, 212.6<br>90, 90, 120 | 54.9, 54.9 210.3<br>90, 90, 120 |
| Total reflections | 140742 (12541) | 80706<sup>a</sup> (8264) | 80598<sup>a</sup> (7871) | 107528 (9723) |
| Unique reflections | 12541 (923) | 4337<sup>a</sup> (420) | 4338<sup>a</sup> (420) | 9723 (719) |
| Multiplicity | 11.2 (11.2) | 18.6 (19.6) | 18.6 (18.7) | 11.1 (9.5) |
| Completeness (%) | 99.8 (100) | 98.74 (100) | 98.94 (100) | 99.8 (100) |
| $I/\sigma(I)$ | 30.7 (1.9) | 41.58 (6.57) | 42.48 (10.86) | 39.45(1.34) |
| $R_{\text{merge}}$ (%)<sup>a</sup> | 0.038 (1.36) | 0.05492 (0.3711) | 0.05491 (0.2373) | 0.035 (1.89) |
| Phasing |  |  |  |  |
| Anomalous resolution (Å) | 2.5 | 3 | 3 | 2.7 |
| Number of sites | 1 | 1 | 1 | 13 |
| Anomalous (%)phasing<br>power<sup>b</sup> | 2.86 | 2.37 | 1.96 | 0.67 |
| Figure of merit acentric | 0.46 | - | - | - |
| Refinement |  |  |  |  |
| Number of reflections<br>used<sup>c</sup> | 7282 (694) | - | - | - |
| $R_{\text{work}}$ (%) | 23 (30) | - | - | - |
| $R_{\text{free}}$ (%)<sup>d</sup> | 24 (44) | - | - | - |
| Number of non-hydrogen atoms in: |  |  |  |  |
| - macromolecule | 1331 | - | - | - |
| - ligand | 1 | - | - | - |
| - solvent | 5 | - | - | - |
| Average B-factor<br>macromolecule (Å<sup>2</sup>) | 102.9 | - | - | - |
| Average B-factor ligand<br>(Å<sup>2</sup>) | 98.3 | - | - | - |
| RMSD bond lengths (Å) | 0.01 | - | - | - |
| RMSD angles (°) | 1.29 | - | - | - |
| Ramachandran favored<br>(%) | 98.6 | - | - | - |
| Ramachandran allowed<br>(%) | 1.4 | - | - | - |
| Rotamer outliers (%) | 0.00 | - | - | - |
| Clashscore | 4.27 | - | - | - |
| PDB code | 8EB5 | - | - | - |

Statistics for the highest-resolution shell are shown in parentheses.
<sup>a</sup> $R_{\text{merge}} = \sum |I_i - \langle I \rangle| / \sum I_i$ , where $I_i$ is the intensity of measured reflection and $\langle I \rangle$ is the mean intensity of all symmetry-related reflections.
<sup>b</sup> Phasing Power is $\langle |F_h(\text{calc})| / \text{phase-integrated lack of closure} \rangle$ where $F_h(\text{calc})$ is the calculated structure factor of the reference scatterers (Zn, P, S).
<sup>c</sup> Friedel's law true.
<sup>d</sup> $R_{\text{free}} = \sum T ||F_{\text{calc}}| - |F_{\text{obs}}|| / \sum F_{\text{obs}}$ , where $T$ is a test dataset of about 3–6% of the total unique reflections randomly chosen and set aside prior to refinement.
RMSD stands for root mean square deviation.

**Table S2.**
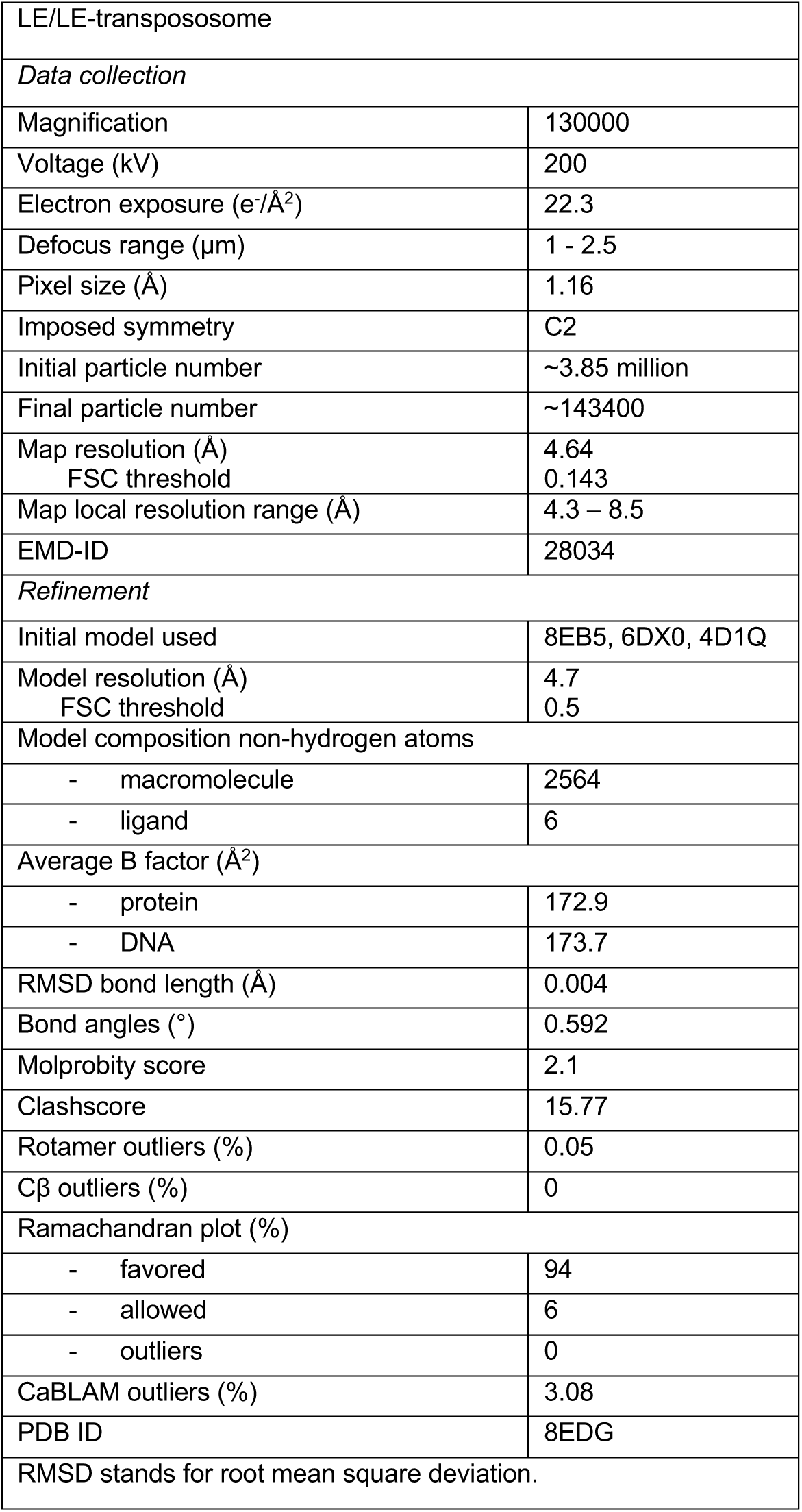
Cryo-EM data collection and refinement statistics

**Table S3.**
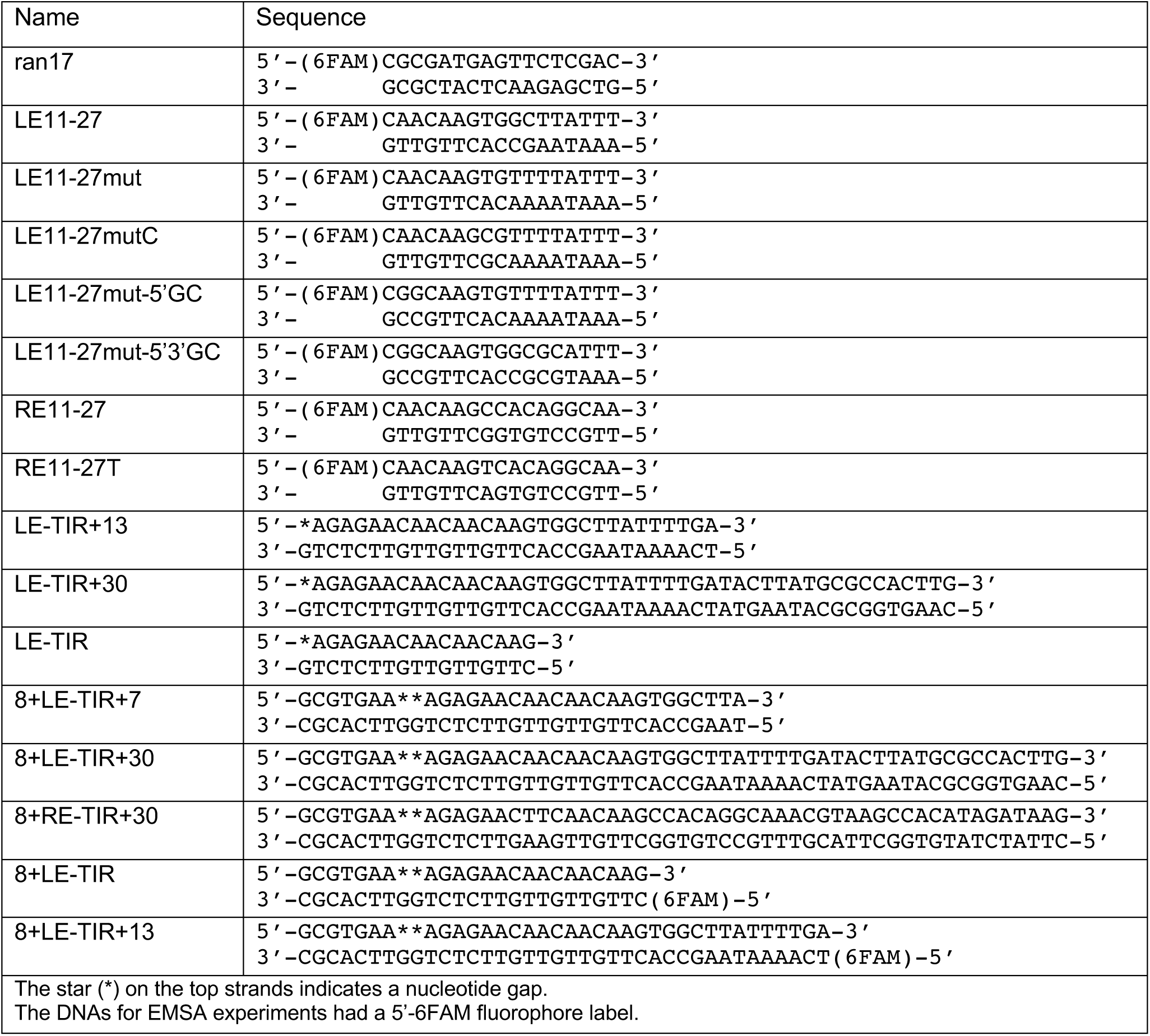
Oligonucleotides used in the study

**Table S4.**
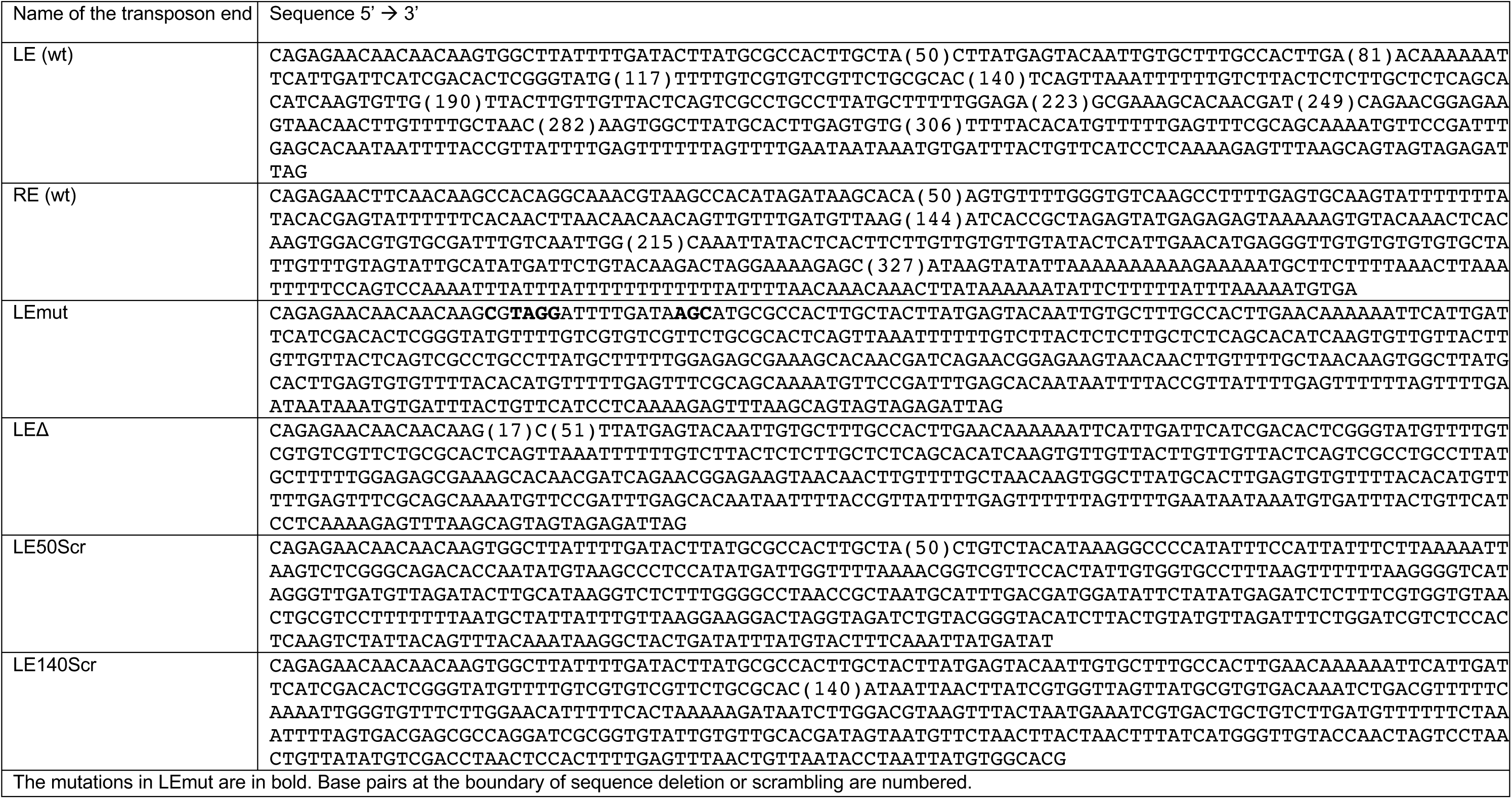
Sequence of the *Hermes* ends of the donor plasmids used in the transposition assay

